# SUC2 sucrose transporter is required for leaf apoplasmic sucrose levels. Consequences for phloem loading strategies

**DOI:** 10.1101/2024.03.20.585851

**Authors:** Françoise Vilaine, Laurence Bill, Rozenn Le Hir, Catherine Bellini, Sylvie Dinant

## Abstract

**Summary**

• The SUC/SUT sucrose transporters belong to a family of active H+/sucrose symporters, with a role of SUC2 in active apoplasmic phloem loading to drive long-distance phloem transport of sucrose in Arabidopsis. However, the cooperation with the symplasmic pathway for phloem loading remains unclear.

• In this study, we explored the consequences of reducing either apoplasmic or symplasmic pathways of phloem loading. We compared a series of lines with modified expression of *SUC2* gene, and we analyzed the effects on plant growth, sugar accumulation in source and sink organs, phloem transport, and gene expression.

• Our data revealed that a modified expression of *SUC2* impacted apoplasmic sucrose levels in source leaves but did not impact phloem transport, as might be expected, while increasing foliar storage of carbohydrates. This response differed from lines in which symplasmic communications between phloem cells was disrupted by the over-expression of a plasmodesmata-associated protein, NHL26.

• Altogether, our studies indicate an unexpected effect of SUC2 for apoplasmic sucrose levels in source leaves, together with SUC1, and suggest a feedback regulation on foliar storage. This data sheds new light on the interplay between symplasmic and apoplasmic pathways for sugar loading and the consequences on leaf water flows.

**Summary statement:** The mechanisms that coordinate apoplasmic and symplasmic loading pathways, and their effects on foliar carbon storage, remain largely unexplored. Surprisingly, the sucrose transporter SUC2 plays a significant role in maintaining sucrose levels in the apoplasm, shedding light on how apoplasmic sugar levels and water flows can interact for phloem loading.

## Introduction

An efficient allocation of photosynthesis products is essential for higher plants to survive as multicellular organisms (Lemoine *et al*., 2013). The phloem regulates the allocation of sugars in the plant, it controls the entry of sugars into the translocation stream (collection phloem), sugar transport from source to sink organs (transport phloem), and delivery to the various competing sink organs (release phloem) (Van Bel, 2003). In most plants, sucrose is the major transport form. Translocated sucrose provides carbon (C) skeletons for primary metabolism, and supplies energy for cellular metabolism (Nunes-Nesi *et al*., 2010). It is an essential signaling molecule for cellular metabolic status (Koch, 2004; Li *et al*., 2021).

Three loading strategies have been described: active loading from the apoplasm, passive diffusion via the symplasm through plasmodesmata (PD), and passive symplasmic transfer followed by polymer trapping (Rennie & Turgeon, 2009). In apoplasmic loaders, sucrose phloem loading involves a passive efflux of sucrose from leaf bundle sheath or phloem parenchyma cells into the phloem cell wall space through SWEET (SUGAR WILL EVENTUALLY BE EXPORTED) sucrose efflux carriers followed by active, proton-coupled import of sucrose into the companion cells (CC) or the sieve elements (SE) via SUC/SUT (SUCROSE TRANSPORTER) proton-sucrose symporters (Braun, 2022). In Arabidopsis, an apoplasmic loader (Haritatos *et al*., 2000b), SWEET11/12 are required for the efflux of sucrose in the apoplasm of the phloem parenchyma cells (Chen *et al*., 2012), and SUC2 is required for its influx into the CC (Gottwald *et al*., 2000). Their action creates a high sucrose concentration in the CC/SE complex that generates the osmotically driven entry of water, generating the mass flow in the SE. Cell-to-cell transport between mesophyll cells and the perivascular cells and trafficking of sucrose between the CC and the SE in the minor veins, where loading takes place, are symplasmic. Plasmodesmata (PD) opening or closing at the interface between CC and SE can also regulate phloem loading. When the phloem plasmodesmal NDR1/HIN1-like protein NHL26 over-accumulates in the PD, it blocks sugar export in Arabidopsis (Vilaine *et al*., 2013). Long-distance transport of sugars from sources to sinks is driven by hydrostatic pressure difference. During its transport to the sinks, sucrose is also released and retrieved continuously into the SE (Hafke *et al*., 2005), its leakage from the SE supplying C to the surrounding tissues (Minchin & Thorpe, 1987). Carbon in excess in the vascular tissues can be stored as starch in the plastids, or as mono or disaccharides in the vacuole, prior to remobilization when sink demand exceeds photosynthetic C supply, as for example during the night.

The coordination of symplasmic or apoplasmic steps and the mechanisms preventing the flow of sucrose back through PD in various cell types (Turgeon, 2006) remain unclear. Both the apoplasmic and symplasmic pathways may contribute to regulation of the photoassimilate flux (Turgeon & Ayre, 2005; Liesche & Patrick, 2017), depending on species, environmental conditions, and developmental stages. For example, in melon, a symplasmic loader, in response to an infection with the cucumber mosaic virus an increased expression of a gene encoding a SUC/SUT sucrose transporter is observed in source leaves, suggesting that active apoplasmic loading takes over the symplasmic transport (Gil *et al*., 2011). However, the mechanisms to coordinate apoplasmic and symplasmic loading pathways, and their consequences on foliar C storage, are still unexplored.

Our previous studies indicated that overexpressing *NHL26* alters sugar allocation in Arabidopsis, with higher sugar accumulation in the rosette and impaired phloem sucrose exudation rate (Vilaine *et al*., 2013). We proposed that the phloem loading of sucrose was blocked in over-expressor lines by the reduced permeability of PD at the interface between CC and SE. However, these lines also showed reduced expression of *SUC2*, and this downregulation may also reduced loading. In this study, we intended to explore the consequences of blocking either apoplasmic or symplasmic loading steps on phloem loading by comparing lines impaired in the expression of either *NHL26* or *SUC2*. Our data demonstrate that SUC2 is required to control the level of sugars in the apoplasm.

## Materials and Methods

### Plant material and growth conditions

The Columbia accession of *Arabidopsis thaliana* (*Col0*) was used in all experiments. The *suc2-4* mutant (SALK_038124) and the partially-complemented line *suc2* x *pGAS:SUC2* (Srivastava *et al*., 2008) were provided by B. Ayre (University of North Texas, USA). The insertional *suc1-3* mutant (Gabi GK139B11) was provided by N. Pourtau (Université de Poitiers, France). It contains a T-DNA insertion in the second intron. The double *suc1 suc2* mutant was obtained by crossing *suc2-4* and *suc1-3*, and homozygous plants were selected with specific primers (Table **S1**). Plants were grown in soil (Tref Substrates) within a growth chamber under long-days (150 µE m^−2^ s^−1^, 16 h light 23°C, 8 h darkness 18°C, 70% humidity). Plants were fertilized with Plant-Prod nutrient solution (Fertil). In these conditions, the floral bud emerged at 28 days after sowing (DAS) in WT plants. The *suc1suc2* plants were grown under short-days (150 µE m^−2^ s^−1^, 10 h light 23°C, 14 h darkness 18°C, 70% humidity), based on previous report indicating that *suc2-4* growth and viability are better under such conditions (Srivastava *et al*., 2009). Projected rosette area (PRA) was measured from pictures and using ImageJ (https://imagej.nih.gov/ij/). Rosette and stem growth rates were measured in the linear part of the growth curve between 1-to-4 weeks or 5-to-9 weeks for the rosette and stem growth, respectively, except for *35S:NHL* and *suc2 stems,* measured between 6-to-9 weeks, and 7-to-10 weeks respectively. The harvest index was calculated as the ratio of seed mass to total aerial dry plant mass measured at harvest. For statistics, Student’s t test was used, with P values <5. 10^-2^ considered significant. Pearson correlations were realized using R software (‘R software, version 3.1.2’).

### Plasmid construction

All constructs were obtained with Gateway technology (Invitrogen). The coding regions of *NHL26* (At5g53730) or *SUC2* (At1g22710) were amplified with specific primers and recombination site-specific sequences. The second step was performed with the primers attB1 and attB2 (Table **S2**), to reconstitute intact attB recombination sites. PCR fragments were introduced into pDONR207 vector (Invitrogen) by BP recombination and transferred by LR recombination into destination vectors (Table **S3**). For overexpression driven by the *CmGAS1* promoter from *Cucumis melo*, the destination binary vector pIPK-pGAS-R1R2-tNOS was obtained by inserting a 3083-bp fragment carrying the promoter region *CmGAS*1 of galactinol synthase, from the pSG3K101 plasmid, provided by Bryan Ayre (Haritatos *et al*., 2000a) into the *Spe*I site of pIPKb001 destination vector (Himmelbach *et al*., 2007). The *amiRNA* to silence *SUC2* gene was designed with WMD3-Web MicroRNA Designer (http://wmd3.weigelworld.org) (Ossowski *et al*.) and inserted into the pRS300 vector. The targeted sequence was 5’-CTACTCGTATATGCAGCGTAT-3’, corresponding to nucleotide positions 778-798 of the *SUC2* ORF, and the *amiRNA* was 5’-TAGATCGCATGACTCAGGCAT-3’ (R complement). The *amiRNA* precursor was transferred into pIPK-pGAS-R1R2-tNOS. For plant transformation, binary vectors were introduced into *Agrobacterium tumefaciens* C58pMP90 (Koncz & Schell, 1986) and plants were transformed by floral dip (Clough & Bent, 1998). Transformants were selected on kanamycin (50 mg/L) or hygromycin (15 mg/L), depending on binary vector. Homozygous T3 seeds were used for phenotypic analyses.

### Phloem sap exudates

Phloem sap exudates were collected by EDTA-facilitated exudation (King & Zeevaart, 1974) with modifications. Leaves were sampled 4 h after the beginning of the light period, after floral transition, at first flower opening. Briefly, the petiole of the 5^th^ or 6^th^ rosette leaf was cut off and recut in exudation buffer (10 mM HEPES buffer pH 7.5, 10 mM EDTA), let 5 min in this buffer then transferred in the collection tube with 80 µl of exudation buffer, for 2 hours exudation in the dark. Exudates of six replicates (one leaf per plant and six plants per genotype) were collected. Soluble sugar and amino acid contents were determined on these samples with no purification step. To determine the contaminations due to leakage from cut cells and from the apoplasm, we used the same protocol, excluding EDTA, which prevents the rapid occlusion of sieve tubes, from the exudation buffer (10 mM HEPES buffer pH 7.5).

### Collect of apoplasmic washing fluids (AWF)

AWF were collected using the infiltration-centrifugation method (Lohaus *et al*., 2001). Plants were grown for 2 months in short-day conditions (150 µE m^−2^ s^−1^, 10 h light 23°C, 14 h darkness 18°C, 70% humidity). Four to six fully developed rosette leaves of six plants were collected, weighed and washed in ice-cold milli-Q water. They were infiltrated with ice-cold milli-Q water containing 0.004% triton X100 by application of a low pressure using a vacuum pump twice for 2 min and wiped dry with tissues, then weighed again. Leaves were then wrapped together, in parafilm and transferred in a syringe (leaf tip pointing downwards). The syringe was placed in a 50 ml Falcon tube and AWF were collected by centrifugation for 20 min at 1000g. The volume of the collected liquid was measured and stored at - 20°C for sugar analysis. For *suc2*, the protocol was slightly modified because leaves were smaller and thicker than WT. About 50 leaves were infiltrated for 10 min, wiped dry with tissues, and placed leaf tip down in a 500µL Eppendorf tube in which a hole has been pierced with a needle. The tube was placed in a 1.5ml tube and the AWF was collected by centrifugation for 20 min at 1000g.

### Soluble sugars, starch, amino acids, total C and N Content

Leaf carbohydrates and amino acids were analyzed using pooled samples from the 3rd, 4th, 7th and 8th rosette leaves collected from the plants utilized for phloem exudate collection. Leaves were harvested 4 h after the onset of the light period and frozen in liquid nitrogen. Soluble sugars and starch were extracted from 50 mg of leaves and quantified using the enzymatic method (Sellami *et al*., 2019). Amino acids leaf content was determined following the Rosen’s method (Rosen, 1957) using the same extracts. Sugars and amino acid contents of phloem exudates and AWF were determined using the same methods, with no hydro-alcoholic extraction. Nitrogen (N) and carbon (C) contents were determined using an elemental analyzer (Thermoflash 2000; Thermo Scientific).

### Protein quantification

Seed and leaf proteins were extracted as described (Lu *et al*., 2020). Two mg of dry seeds or 50 mg of leaf tissue (5^th^ or 6^th^ leaf) were ground in the extraction buffer (100 mM Tris-HCl (pH8), 0.1% [w/v] SDS, 10% [v/v] glycerol] and 2% [v:v] 2-mercaptoethanol). The samples were centrifuged at 14,000g at 4°C for 10 min, and the supernatants were collected. Proteins were quantified using Bio-Rad Protein Assay.

### Lipid quantification in seeds

Lipid were extracted and quantified from seeds as described (Reiser *et al*., 2004). First, 0.1 g of air-dried seeds was ground in liquid nitrogen, 1.5 mL isopropanol was added, the resulting extract was transferred into a 1.5-mL reaction tube and incubated with agitation for 12 h at 4°C at 100 rpm. Subsequently, the mixture was centrifuged at 12,000 g for 10 min. The supernatant was transferred into a 1.5-mL tube, then incubated at 60°C overnight to allow the evaporation of isopropanol. Total lipid was quantified gravimetrically.

### RNA Isolation and Q-RT-PCR

A portion of the leaf material collected for sugar and starch quantification was utilized for RNA extraction. Total RNA was isolated from frozen tissue using TRIZOL (ThermoFisher Scientific). Reverse transcription was conducted with 1 µg total RNA with the Superscript II enzyme (Invitrogen), after DNase treatment (Invitrogen). The primers employed for Q-PCR amplification are listed in Table **S4**. qRT-PCR was carried out with the MESA GREEN MasterMix Plus for SYBR assay, following the manufacturer’s instructions (Eurogentec). Amplification was performed with 1 µL of a 1:10 or 1:20 dilution of cDNA in a total volume of 10 µL: 5 min at 95°C, followed by 40 cycles of 95°C for 5 s, 55°C for 15 s, and 72°C for 40 s, in an Eppendorf Realplex2 MasterCycler (Eppendorf SARL). Two reference genes, *TIP41* (At4g34270) and *APT1(At1g27450),* were utilized, yielding comparable results. The data are presented as percentages of *TIP41* expression. Heat maps were generated after normalization by the mean value of gene expression in WT plants and visualized on Genesis 1.8.1 software on a log_2_ scale.

## Results

### Impairing the symplasmic pathway by an over-expression of NHL26

The *p35S::NHL26* line (hereafter referred to as *35S:NHL*) carries the coding region of *NHL26* under the 35S promoter (Vilaine *et al*., 2013) (Fig. **S1a**). The *NHL26* transcript accumulated substantially in p35S:NHL lines (300 to 800 times normal levels). We created new transgenic lines in which *NHL26* over-expression was targeted to the CC of the minor veins in mature leaves, using the *CmGAS1* promoter from *Cucumis melo* (Haritatos *et al*., 2000a) (Fig. **S1a**). These *pCmGAS::NL26* lines (hereafter called *GAS:NHL*) present an upregulation of *NHL26* (Fig. **S1b,c**) and showed reduced growth and an increased accumulation of soluble sugars in source leaves compared to wild-type plants (Fig. **S1d**).

Line *GAS:NHL#12, in which NHL26* transcript accumulated significantly (100 times normal level) was selected for detailed characterization and comparison with *35S:NHL* line (Fig. **1**). Both lines showed a reduced rosette and stem growth compared to the wild-type, partly due to the delayed flowering in *35S:NHL* line (Fig. **1a-g**), associated with an increased accumulation of soluble sugars and amino acids in the rosette (Fig. 1**h,i**). Starch content, bolting time, and harvest index were unchanged in *GAS:NHL#12* plants compared to the wild-type, while they were reduced in *35S:NHL* plants (Fig. **1j-l**).

**Fig. 1.**
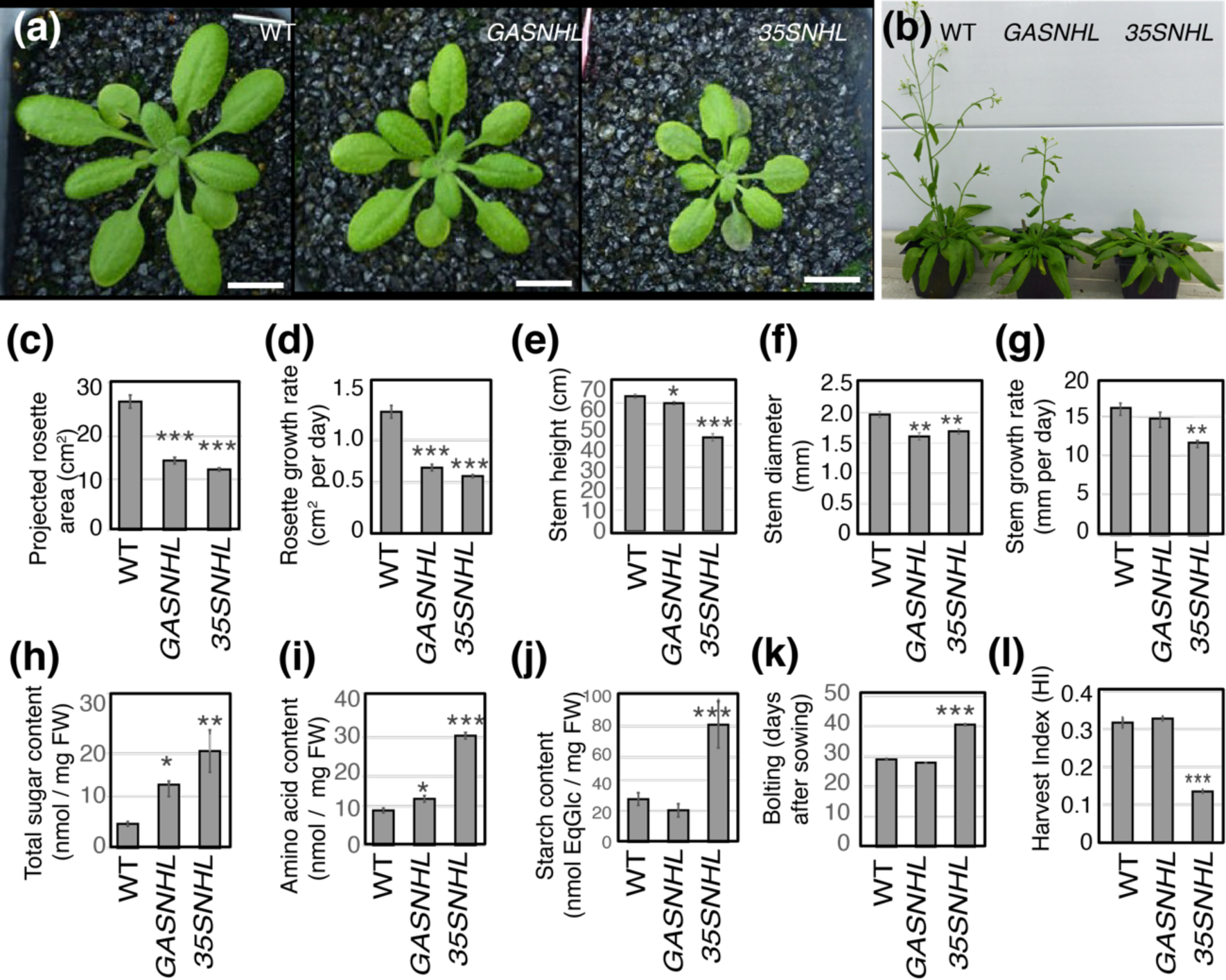
Phenotype of the *GAS:NHL* and *35S:NHL* plants. **(a):** Phenotype of 3 weeks-old plants. **(b):** Phenotype of 6 weeks-old plants. **(c):** Projected rosette area (PRA) of 4 weeks-old plants (cm^2^). **(d):** Rosette growth rate between 7 and 21 days after sowing (DAS) (cm^2^ per day). **(e):** Floral stem height of 10 weeks-old plants (cm). **(f):** Floral stem diameter of 10 weeks-old plants (mm). **(g):** Floral stem growth rate between 35 and 56 DAS (mm per day). **(h):** Total soluble sugar content (sucrose, glucose and fructose) in nmoles per mg of fresh weight (FW) in rosette leaves. **(i):** Total amino acids content in nmoles per mg of FW in rosette leaves. **(j):** Starch content in nmoles EqGlucose per mg of FW in rosette leaves. **(k):** Bolting time (DAS). **(l)**: Harvest index (HI). Bar plots and error bars represent the mean and *se* (*n* = 6). Asterisks indicate significant differences compared to control plants (* p < 0.05; ** p < 0.01; *** p < 0.001).

Compared to the wild-type, seed protein and lipid contents and 1000-seed-weight were not altered in the *GAS:NHL#12* plants unlike in the *35S:NHL* plants (Fig. **2a-c**). No modification of sugar and starch contents was observed (Fig. **2d,e**). The percentage of N was increased and that of C was reduced in the *35S:NHL* plants (Fig. **2f,g**). This resulted in a significantly reduced C/N ratio in *35S:NHL* seeds (Fig. **2h**). The C/N ratio was also reduced in *GAS:NHL#12 seeds,* due to slight -although not significant - variations of seed C and N percentage compared to the wild-type (Fig. **2f,g**). The data indicate that over-expression of *NHL26* in the CC of the minor veins of source leaves reduces plant biomass and increases rosette sugar content. However, the impact is less pronounced compared to what is observed in *35S:NHL* plants.

**Fig. 2.**
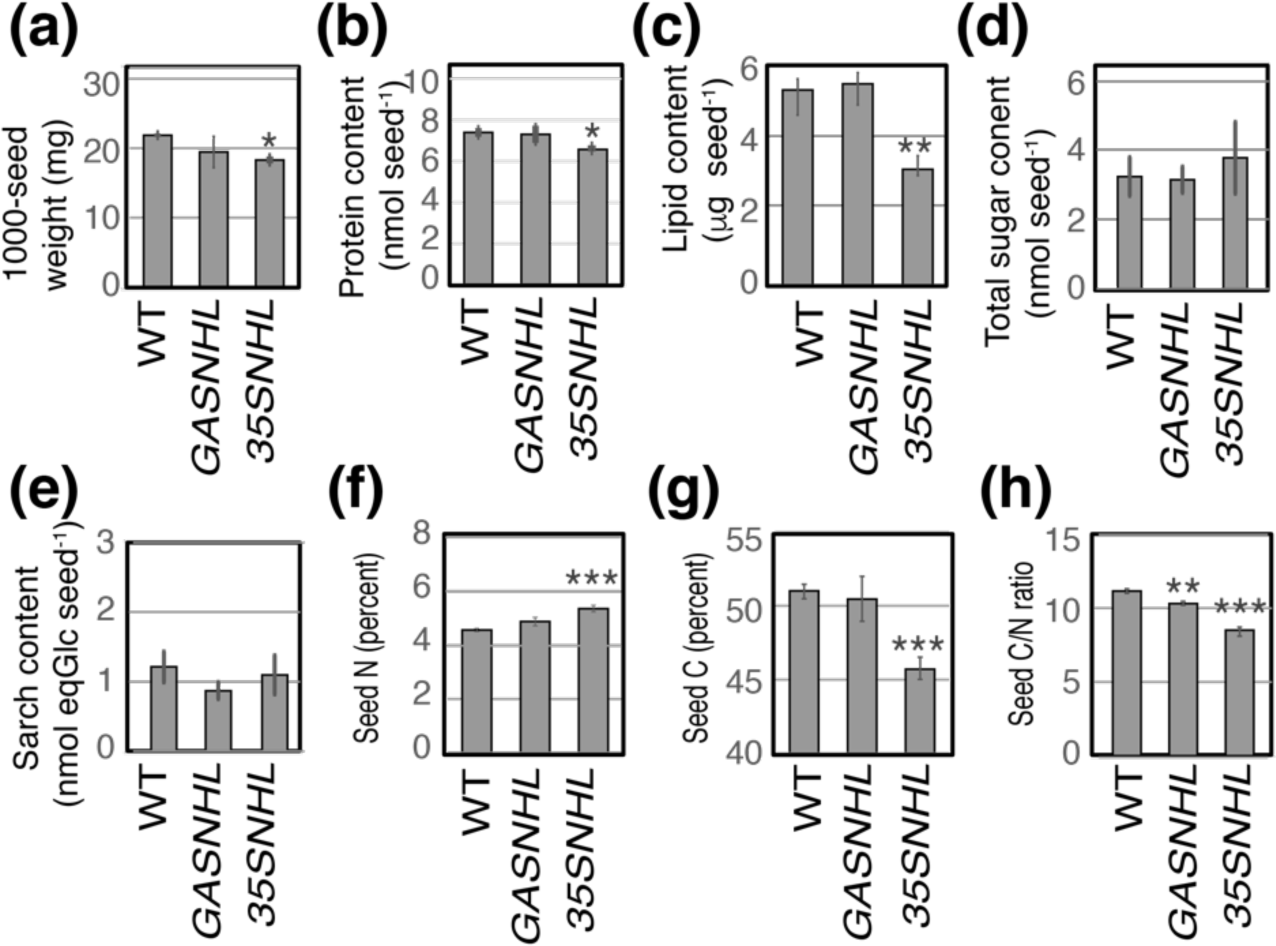
Seed phenotype of the *GAS:NHL* and *35S:NHL* plants. **(a):** Weight of 1000-seeds (mg). **(b):** Protein content (nmoles/seed). **(c):** Lipid content (µg/seed). **(d):** Total sugar content (sucrose, glucose and fructose) (nmoles/seed). **(e):** Starch content (nmoles EqGlucose /seed). **(f):** Percentage of N in seeds. **(g):** Percentage of C in the seeds. **(h):** C/N ratio in seeds. Bar plots and error bars represent the mean and *se* (*n* = 6). Asterisks indicate significant differences compared to control plants (* p < 0.05; ** p < 0.01; *** p < 0.001).

Overall, when *NHL26* is overexpressed in minor veins only, i.e. when PD permeability is potentially modified in the collection phloem, the effects observed are milder then when *NHL26* is ubiquitously overexpressed. Although there is no repression of *SUC2* expression in the *GAS:NHL#12* line, unlike in the *35S:NHL* line (Fig. **S1e,f**), the phenotypes can still result from impaired sucrose loading as proposed for *35S:NHL* plants (Vilaine *et al*., 2013), associated with negative feedback on *SUC2* expression in the minor veins.

### Impairing the apoplasmic pathway by a reduced expression of SUC2

To test the hypothesis that a reduced expression of *SUC2* in the minor veins could affect the overall growth, we analyzed lines in which *SUC2* was either knocked-out (*suc2* mutant), or partially restored in the minor veins of *suc2* mutant (*suc2* x *pGAS:SUC2* line, hereafter referred to as *suc2pC*) (Srivastava *et al*., 2008), or specifically silenced in the minor veins of source leaves, creating new lines expressing an *amiRNA* targeting *SUC2* and driven by the *CmGAS* promoter (Fig. **S2a**). These lines (hereafter referred to as *miSUC*) have a significantly reduced *SUC2* transcript amount (Fig. **S2b**), associated with a reduced rosette and floral stem growth (Fig. **S2c,d**) and an increased sugar accumulation in source leaves compared with the wild-type (Fig. **S2e**). These phenotypes were more severe as *SUC2* expression decreased (Fig. **S2f**). Two representative lines (*miSUC*#4 and #12), *suc2* and *suc2pC* were further characterized (Fig. **3**). A reduction in plant growth was observed at both vegetative and reproductive stages, correlating with the overall decrease in *SUC2* expression (Fig. **3a,b**), while the tissues in which *SUC2* expression is altered are distinct.

**Fig. 3.**
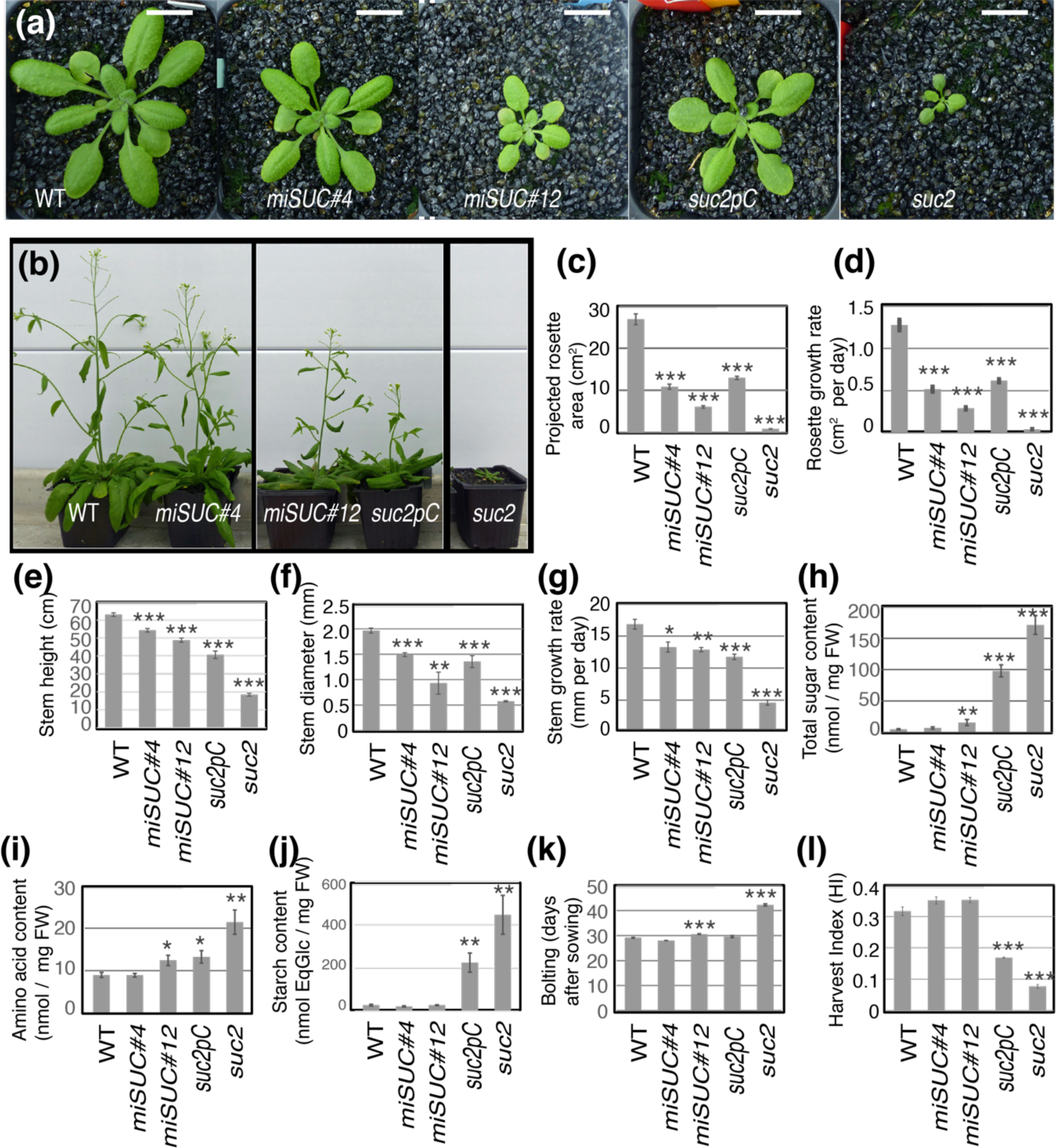
Phenotype of the *SUC* lines. **(a):** Phenotype of of 3 weeks-old plants. **(b):** Phenotype of 6 weeks-old plants. **(c):** Projected rosette area (PRA) of 4 weeks-old plants (cm^2^). **(d):** Rosette growth rate between 7 and 21 days after sowing (DAS) (cm^2^ per day). **(e):** Floral stem height of 10 weeks-old plants (cm). **(f):** Floral stem diameter of 10 weeks-old plants (mm). **(g):** Floral stem growth rate between 35 and 56 DAS (mm per day). **(h):** Total soluble sugar content (sucrose, glucose and fructose) in nmoles per meg of fresh weight (FW) in rosette leaves. **(i):** Total amino acids content in nmoles per mg of FW in rosette leaves. **(j):** Starch content in nmoles EqGlucose per mg of FW in rosette leaves. **(l):** Bolting time (in days). (**l**): Harvest Index (HI). Bar plots and error bars represent the mean and *se* (*n* = 6). Asterisks indicate significant differences compared to control plants (* p < 0.05; ** p < 0.01; *** p < 0.001)

When *SUC2* is knock-out, rosette and stem growth were reduced (Fig. **3c-g**), with a dramatic increase in the accumulation of soluble sugars, amino acids and starch (Fig. **3h-j**) and a delay in flowering and a reduced harvest index (Fig. **3k,l**), consistent with previous reports (Srivastava *et al*., 2008). In *suc2pC,* where *SUC2* expression is restored in minor veins, we observed partial complementation of plant growth (Fig. **3c-g**), associated with less spectacular levels of leaf sugars, amino acids and starch, although they remained high compared to WT (Fig. **3h-j**). In *miSUC*#4, *miSUC*#12, where *SUC2* expression is silenced in minor veins, the growth phenotypes were similar to those of *suc2pC* (Fig. **3c-g**), whereas the effects on loading and retrieval are theoretically opposite. By contrast, soluble sugar levels were barely affected in *miSUC*#12 with no effect in *miSUC*#4 (Fig. **3h**), and starch levels were unaffected compared with WT (Fig. **3j**). Regarding seed weight and quality, which are severely impaired in *suc2*, we did not observe any complementation of seed weight, protein and lipid content, seed N and C content in the *suc2pC* plants (Fig. **4a-h**), although sugar and starch content were unchanged compared with WT (Fig. **4d,e**), In contrast, in *miSUC*#4, *miSUC*#12, none of these traits were altered (Fig. **4a-h**).

**Fig. 4.**
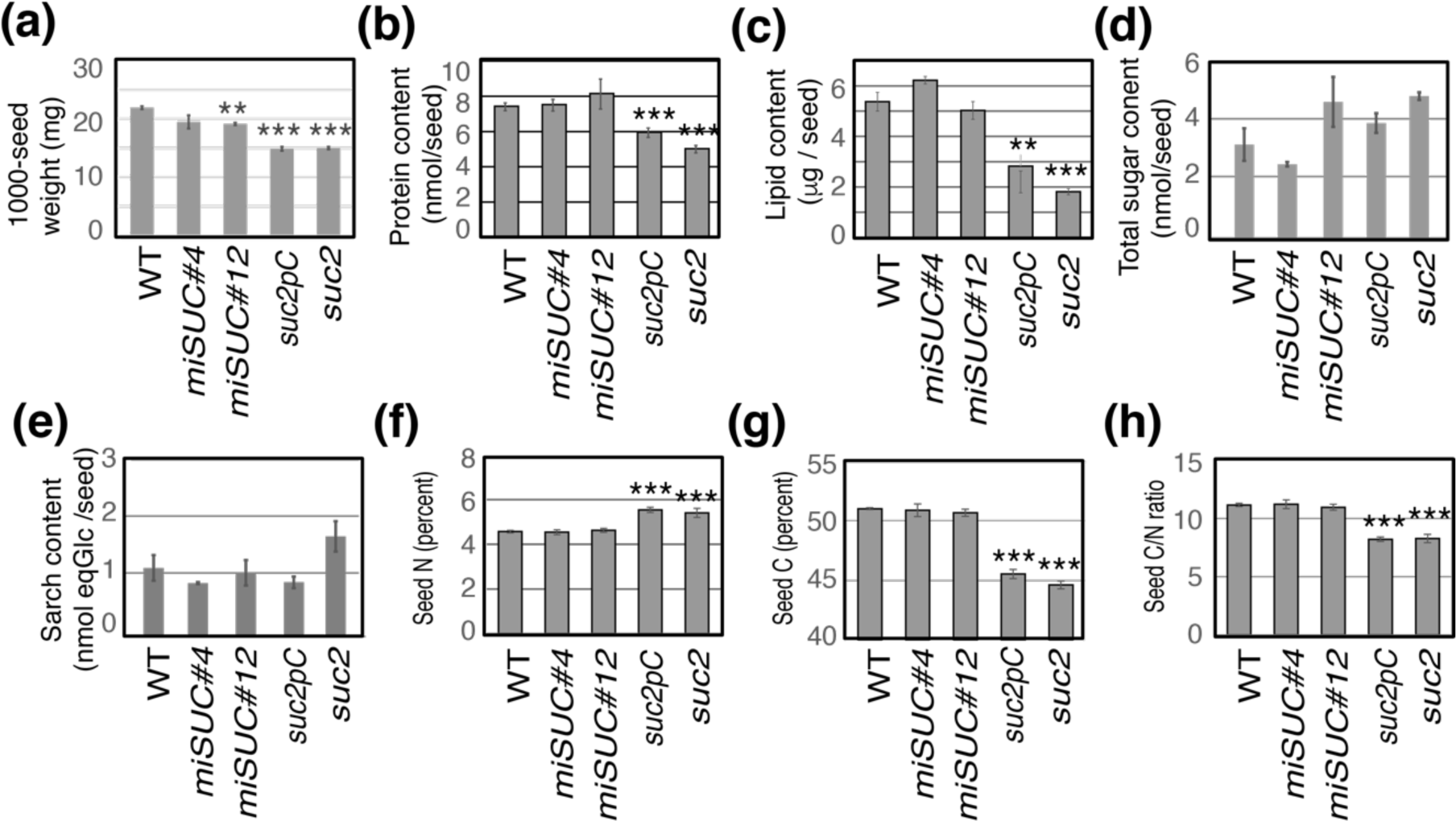
Seed phenotype of the *SUC* lines. **(a):** Weight of 1000-seeds (mg). **(b):** Protein content (nmole/seed). **(c):** Lipid content (µg/seed). **(d):** Total sugar content (sucrose, glucose and fructose) (nmole/seed). **(e):** Starch content (nmol EqGlucose /seed). **(f):** Percentage of N in seeds. **(g):** Percentage of C in the seeds. **(h):** C/N ratio in seeds. Bar plots and error bars represent the mean and *se* (*n* = 6). Asterisks indicate significant differences compared to control plants (* p < 0.05; ** p < 0.01; *** p < 0.001).

### Impacts of a modification of SUC2 or NHL26 expression on phloem sugar exudation

The reduced growth in the six genotypes compared with the WT, associated with an excess of sugars in the rosette leaves in *GAS:NHL, 35S:NHL, suc2pC and suc2 plants,* is consistent with an alteration of phloem transport. To test this hypothesis, we collected phloem exudates using EDTA-facilitated exudation method, then measured sucrose and hexoses to calculate sugar exudation rates from cut leaf petioles. In the *35S:NHL* plants, we observed a reduced sucrose exudation rate compared with wild-type plants (Fig. **5a**), which is consistent with the initial study of *35S:NHL* lines (Vilaine *et al*., 2013) that concluded that over-accumulation of NHL26 protein leads to the closure PD and blocks sugar loading. The same tendency was observed in the *GAS:NHL* plants, although it was not significant. In *suc2pC* plants, where SUC2 expression is restored in collection phloem, the rate of sucrose exudation was not reduced compared to wild-type plants, consistent with complementation of phloem loading. Surprisingly, when *SUC2* was silenced in the minor veins only (*miSUC#4 and #12* plants), where we expect an impact on phloem loading, the rate of sucrose exudation was not different than that of the wild-type (Fig. **5a**). In *suc2* plants, the sucrose exudation rate was dramatically increased compared to wild-type plants with more glucose and fructose in the exudates as well (Fig. **S4**). The simplest hypothesis to explain the data would be that the increase in sucrose and hexoses results from contamination due to increased leakage from cut petioles.

**Fig. 5.**
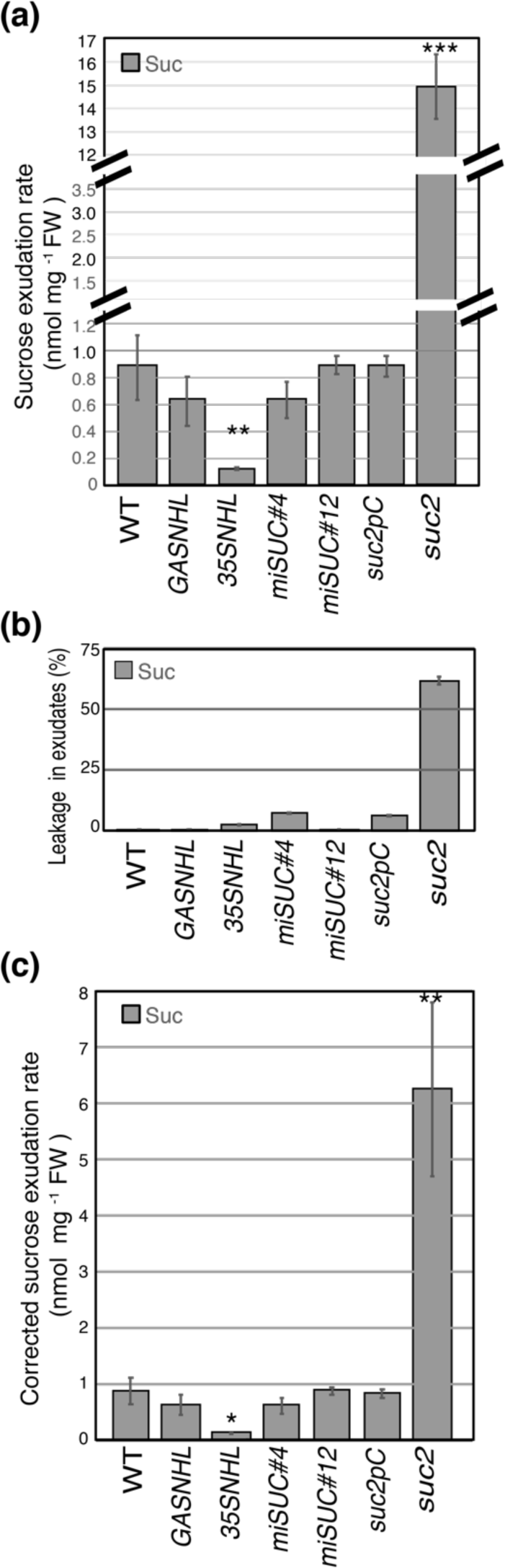
Rate of phloem sucrose exudation in *NHL* and *SUC* plants. **(a):** Sucrose exudation rate was measured on the phloem exudate collected by EDTA-facilitated exudation of one rosette leaf per plant (5^th^ or 6^th^ leaf) for 2 hours of exudation. **(b):** Sucrose leakage was measured on the exudate on a second rosette leaf of the same plant (5^th^ or 6^th^ leaf) collected using the same method but omitting EDTA. **(c):** Corrected exudation rate of sucrose, calculated by subtracting sucrose leakage value from the exudation rate. Bar plots and error bars represent the mean and *se* (*n* = 6). Asterisks indicate significant differences compared to control plants (* p < 0.05; ** p < 0.01; *** p < 0.001). FW: fresh weight.

In the EDTA-facilitated exudation protocol, EDTA chelates Ca^2+^ ions that would otherwise participate in phloem sealing processes (King & Zeevaart, 1974). The omission of EDTA from the exudation buffer can reveal sugar leakage by a route other than phloem translocation. We measured sucrose in petiole exudates obtained after excluding EDTA from the exudation buffer (Fig. **5b**). The data indicate that about 50% of the sugars present in the exudate from *suc2* leaves obtained in the presence of EDTA were also present in exudates without EDTA, revealing high levels of leakage occurring in *suc2* leaves. Such contamination was not observed with the other genotypes, except for the *35S:NHL* plants with more hexose leakage than the wild-type (Fig. **S3a**).

To take into account leakage, we corrected the rate of sucrose exudation, by subtracting the leakage values measured in the exudates collected without EDTA from the values of sucrose measured in the exudates collected with EDTA (Fig. **5c**). As expected, the corrected exudation rate of sucrose remained reduced in *35S:NHL* plants, which is consistent with earlier report (Vilaine *et al*., 2013). It was not significantly different in the *GAS:NHL*, *miSUC#4 and miSUC#12* plants compared to the wild-type. However, it was significantly higher in *suc2* plants (6-fold increase). The corrected values for hexoses were also higher in *suc2* plants than in all other lines (Fig. **S3b-e**), revealing in *suc2* an increase in the proportions of hexoses in the phloem exudates (Fig. **S3f**). Interestingly, *SUC2* expression in minor veins of the *suc2* mutant (*suc2pC plants*) totally complemented the wild-type sucrose exudation rate (Fig. **5a**; Fig. **S4a-c**).

### Impacts of a modification of SUC2 or NHL26 expression on apoplasmic washing fluids (AWF)

The data suggested an excess of sugars in the apoplasm. We quantified sugar contents in the AWF collected from rosette leaves. The results revealed elevated sucrose, glucose, and fructose levels in the AWF from *suc2* mutant compared to wild-type plants (Fig. **6a-c**). Sugar levels were also higher, albeit to a lesser extent, in the AWF from all the other lines compared to wild-type plants (Fig. **6a-c**). Notably, two-fold change was observed in AWF sucrose levels in *GAS:NHL* and *35S:NHL,* about four-fold change in *miSUC#4* and *miSUC#12* and over 20-fold change in the *suc2pC* line. These fold changes observed in the AWF surpassed those noted in the corrected sucrose exudation rates (Fig. **S4a**). The data indicate that the altered expression of *SUC2* or *NHL26* in either the loading or transport phloem was associated with changes in apoplasmic sugar levels.

**Fig. 6.**
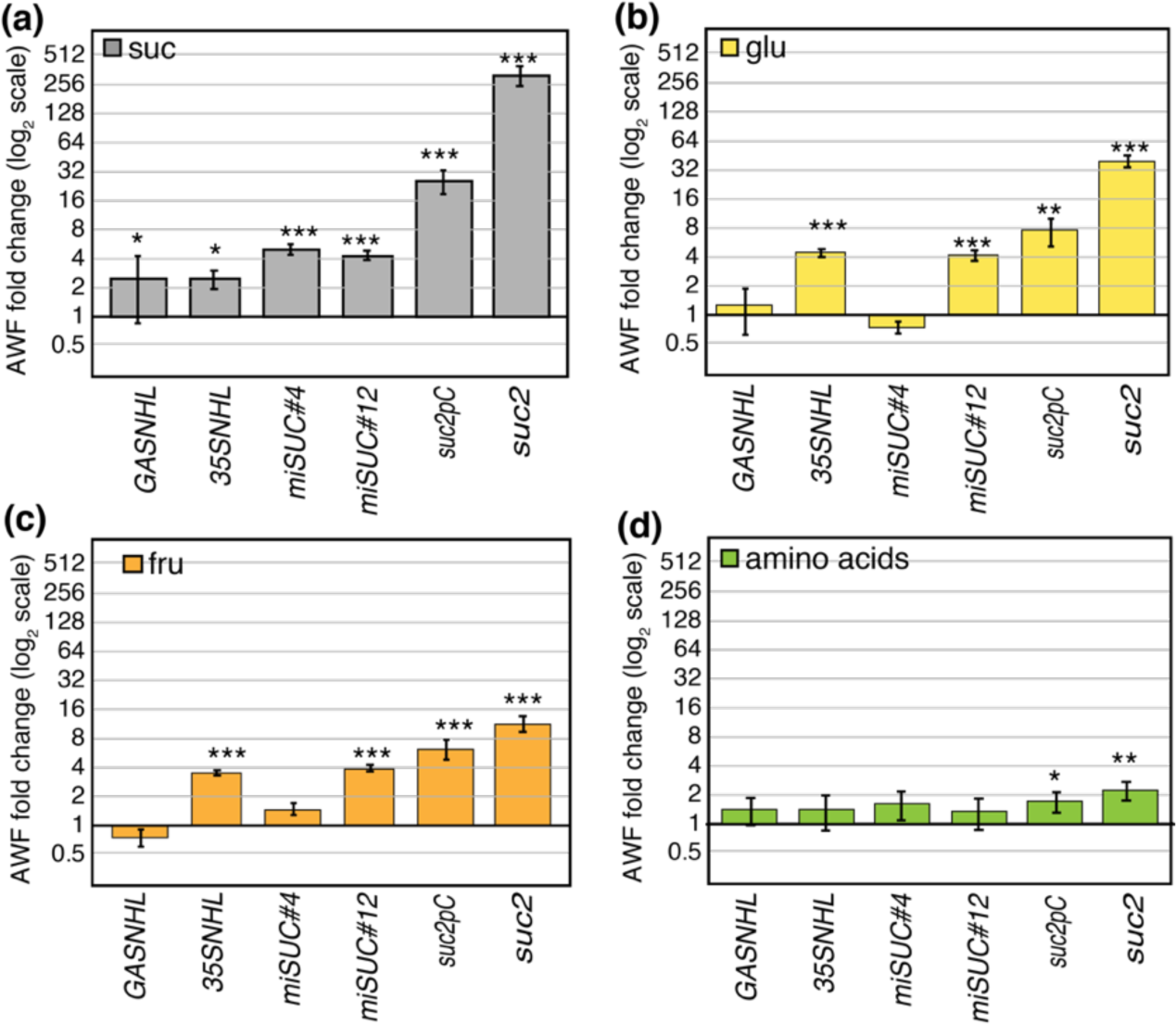
Sugars and amino acids in the apoplasmic washing fluids in *NHL* and *SUC2* plants. **(a)** Sucrose, (**b**) glucose, (**c**) fructose and (**d**) amino acids contents (measured in nmoles per mg of fresh weight) in the leaf apoplasmic washing fluids (AWF), and expressed as fold change compared to WT.

We also measured amino acid contents in the AWF. The results revealed a modest increase in amino acid levels in the AWF from the *suc2* and *suc2pC* plants, reaching a maximum two-fold change (Fig. **6d**) compared to the wild-type. These effects were of a similar magnitude to those observed in corrected phloem exudation rates (2-to 6-fold change compared to the wild-type) (Fig. **S4d**). The data suggest that the altered expression of *SUC2* or *NHL26* has limited impact on amino acid levels compared to the pronounced effects on sugar levels.

### Transcriptional responses to impaired phloem sugar loading

The important modifications in sugar contents observed in phloem exudates and AWF may lead to defects in sugar homeostasis. We analyzed the expression of a subset of genes (Table **S5**), coding for sugar transporters and expressed in different leaf tissues (Fig. **7a**). It includes genes from the *SUC/SUT*, *SWEET* and *STP* (*SUGAR TRANSPORTER PROTEIN*) families coding for disaccharide and monosaccharide transporters (*SUC1-5*, *SWEET16-17*, *SWEET11-12*, *STP1/13*). It includes genes coding glycolytic enzymes (fructokinases FRK1, 2, 3, 6, 7; cell wall invertases cwINV 1, 3 6; cytosolic invertases cINV1-2, and vacuolar invertases vINV1-2), sugar signaling components (hexokinases *HXK1*, *HXK2*, and trehalose phosphate synthase *TPS5*), and other genes: *GPT2* which encodes the plastidial sugar translocator2, *APL3* and *APL4* that encode ADP-Glc pyrophosphorylase large subunits for starch synthesis, *PAP1* which encodes the PRODUCTION OF ANTHOCYANIN PIGMENT1 transcription factor and *GSTF12* which encodes a GLUTHATIONE-S TRANSFERASE12 involved in anthocyanin trafficking. *RRTF1* (REDOX RESPONSIVE TRANSCRIPTION FACTOR 1), *XIP1* (*XYLEM INTERMIXED WITH PHLOEM 1*) and *TRAF-like1 (TUMOR NECROSIS FACTOR RECEPTOR ASSOCIATED FACTOR),* which encodes signaling regulators acting upstream of primary metabolism. Two photosynthesis genes (*LHCB1* and *RBCS*) were also included.

**Fig. 7.**
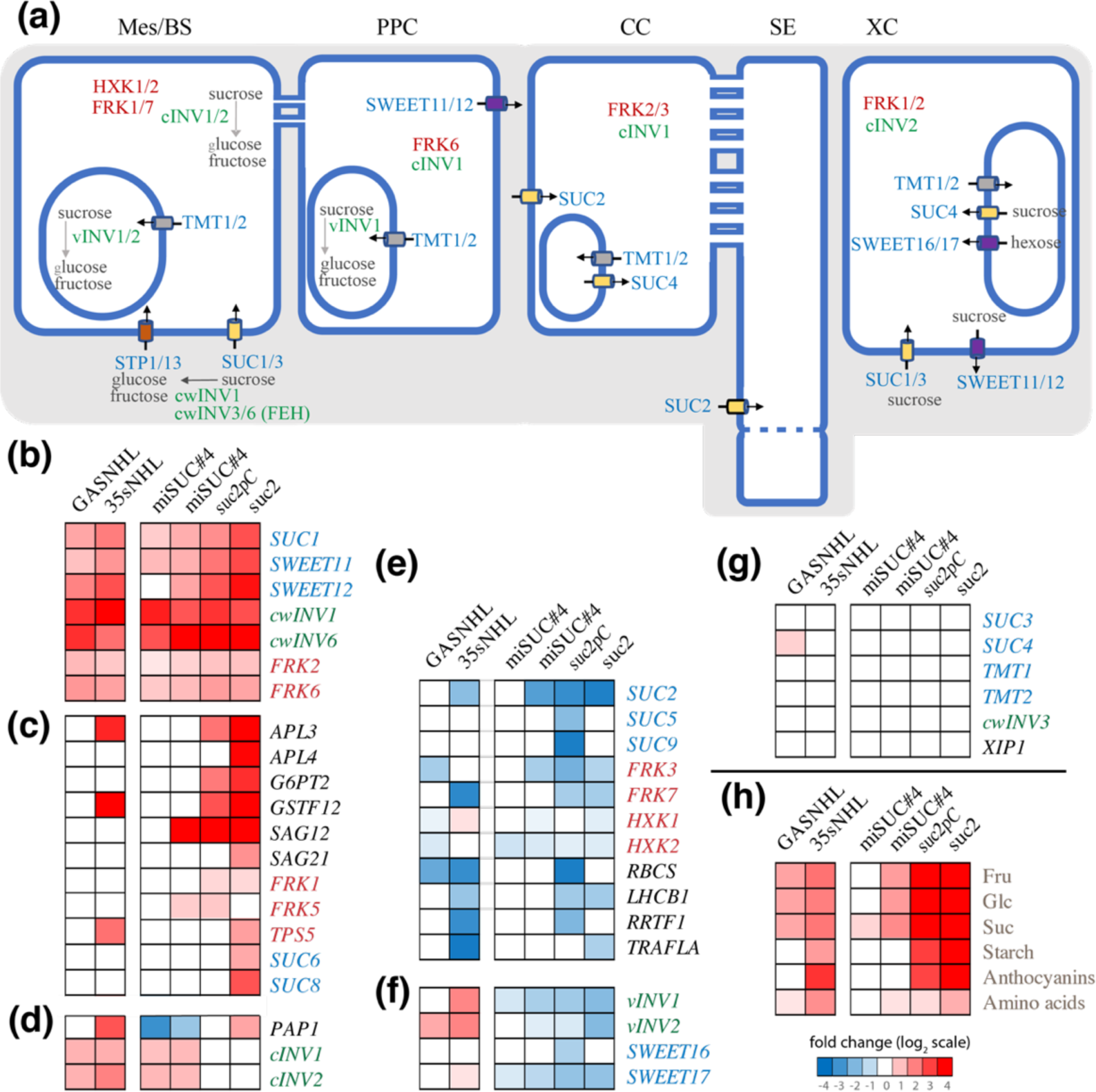
Fold change in transcript amounts in *NHL* and *SUC* plants. (**a**): Cell type specific expression of the selected genes (modified from Xu & Liesche 2021) (Xu & Liesche, 2021). In blue letters: genes coding sugar transporters, in green, genes coding invertases, in red, genes coding FRK, HXK or TPS. In black miscellaneous. Mes: mesophyll, BS: bundle sheat, PPC: phloem parnehcyma cells, CC: companion cells, SE: sieve elements, XC: xylem cells. (**b**) to (**g**): Heat map showing fold changes in relative transcript amount for selected genes in the *NHL* and *SUC* lines compared to WT. (**b**) **(c) and (d):** genes showing an upregulation in *NHL* and/or *SUC* lines. (**e**): genes showing a downregulation in *NHL* and/or *SUC* lines. (**f**): genes showing opposite response in *NHL* or *SUC2* lines. (**g**) genes showing no significant changes in *NHL* and *SUC* lines. (**h**) Heat map showing fold changes in soluble sugars, starch, anthocyanins and amino acids content in rosetted. Fold change are shown in a log_2_ scale, with blue indicating significant lower values in *NHL* or *SUC* lines compared to WT, and in red indicating higher values (*p* < 0.05, *n* = 4-6).

We observed a higher accumulation of *SUC1*, *SWEET11*, *SWEET12*, *CwINV1*, *CwINV6*, *FRK2,* and *FRK6* transcripts in the lines with modified expression of *SUC2* or *NHL26* compared to wild-type plants (Fig. **7b**). A higher transcript amount was observed in the plants showing the highest sugar content for the genes involved in starch biosynthesis (*APL3*, *APL4*, *G6PT2*), glycolysis or sugar signaling (*FRK1*, *FRK5* et *TPS5*), anthocyanin biosynthesis (*GSTF12*, *PAP1*) and *SUC6* and *SUC8* (Fig. **7c-d**). Interestingly, the expression of some of these genes was correlated to leaf sucrose content (Table **S6**). The *SUC3*, *SUC5,* and *SUC9* transcript amounts were either unchanged or slightly reduced in the lines with altered expression of *SUC2* or *NHL26* compared to wild-type plants (Fig. **7e,f**). Interestingly, the *SWEET17* (coding tonoplastic facilitator) transcript amount was reduced in the *miSUC, suc2pC* and *suc2* plants, unlike GAS:*NHL* and *335S:NHL* plants, in which there was either no change or slight upregulation compared to WT. A similar response was observed for *vINV1* and *vINV2,* coding vacuolar invertases. No variations were observed in the accumulation of *TMT1/TST1* and *TMT2/TST2* transcripts, which code tonoplastic transporters. The *cINV1 and cINV2 (*coding cytosolic invertases) transcript amounts were higher in GAS:*NHL, 335S:NHL and miSUC2* plants, unlike *suc2pC* and *suc2* plants, in which there was no change compared to WT. Finally, the *ERDL6 (*coding tonoplastic glucose exporter) transcript amount was higher in *GAS:NHL, 335S:NHL and miSUC2#4* plants compared to WT.

### Additive effects of suc1 and suc2 mutations on plant growth

There was an upregulation of *SUC1* expression was observed in *NHL* and *SUC* lines, suggesting a potential functional complementation of *SUC2* by *SUC1*. Both SUC1 and SUC2 are low-affinity sucrose-transporters with similar sucrose transport activity (Sauer & Stolz, 1994). *SUC1* complements *SUC2* in the *suc2* mutant when expressed from the *SUC2* promoter (Wippel & Sauer, 2012). The function of *SUC1* during the vegetative stage is unclear since *suc1* has a wild-type phenotype for rosette growth (Sivitz *et al*., 2008). In investigating the possibility of complementation of the *suc2* mutation through the overexpression of *SUC1*, we analyzed the growth phenotype of the *suc1suc2* double mutant (Fig. **S5**). When grown under short-days, the *suc2* plants exhibited reduced growth compared to *suc1* plants, accompanied by anthocyanin accumulation. Notably, the *suc1suc2* double mutant was smaller than *suc2*, indicating an additive effect with respect to plant growth (Fig. **S5**).

## Discussion

SUC/SUT transporters have been identified in both symplasmic and apoplasmic loaders (Julius *et al*., 2017). There is a potential for the regulation of those pathways to be coordinated, which could utilize SUC/SUT transporters. In Arabidopsis, which, according to Gamalei’s definition (Gamalei, 1989), is a type 1-2a apoplasmic loader (Haritatos *et al*., 2000b), there are PD at the interface between phloem parenchyma cells and companion cells (PPC/CC). SUC2/SUT1 provides influx of sucrose into the CC (Truernit & Sauer, 1995; Stadler & Sauer, 1996). Our current understanding is that SUC2 contributes to phloem loading by increasing in the CC/SE complex the osmotic potential that drives water flow into the SE (Gottwald *et al*., 2000). SUC2 has also been proposed to retrieve the sucrose leaking from the SE back to the transport phloem (Gould *et al*., 2012). Here, we investigated the possibility of an interplay of *SUC2*-dependent apoplasmic and symplasmic pathways for phloem loading. In *OEX:NHL26* lines, we have postulated that the abnormal buildup of NHL26 at the PDs alters the permeability of PDs between CC and SE (Vilaine *et al*., 2013), hindering the symplasmic exchange between CC and SE and consequently decreasing phloem loading. Sucrose accumulation in the CC was proposed to trigger negative feedback regulation on the SUC2-dependent sucrose influx, impairing the apoplasmic pathway and phloem loading. In the new series of lines with a modified expression of *NHL26* or *SUC2*, we observed elevated accumulation of sugars, starch, and amino acids in the source leaves, reduced rosette and floral stem growth and reduced seed production. A reduced C/N ratio in the seeds was also observed, revealing reduced carbon and lipid contents. Our findings are consistent with the ^14^C labeling studies with *suc2* KO mutants (Gottwald *et al*., 2000), where carbon allocation from sources to sinks was reduced.

### An unsuspected role for SUC2 and SUC1 in the regulation of sucrose levels in the leaf apoplasm

Surprisingly, despite the downregulation of *SUC2* expression in the minor veins’ CC, the sucrose exudation rate, which serves as a proxy for phloem loading, remains unimpaired in the *miSUC* lines. This suggests that *SUC2* expression in the minor veins’ CC may not be essential for maintaining phloem flow. Intriguingly, we observed a concurrent increase in leaf apoplasmic sucrose levels, correlating with the downregulation of *SUC2*. These alterations were also observed in the *suc2pC* line, providing *SUC2* function in the minor veins but defective in *SUC2*-dependent sucrose retrieval in the transport phloem. Our findings suggest that *SUC2* expression in either the collection phloem or the transport phloem affects leaf apoplasmic sucrose levels, and indicate it is not needed to provide phloem mass flow (Fig. **8a-c**).

**Fig. 8.**
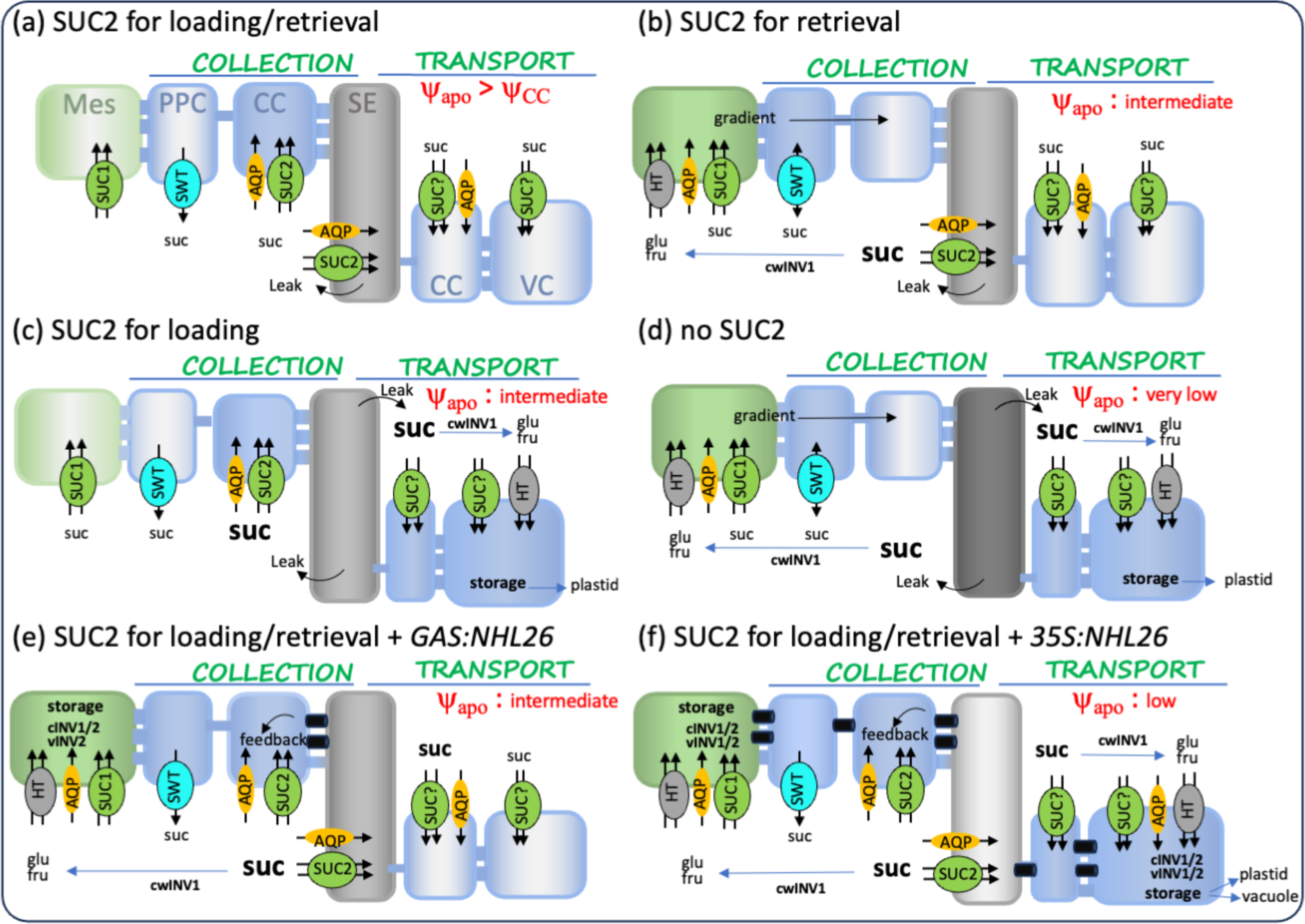
Proposed model of the roles of SUC1 and SUC2 in regulating apoplasmic sucrose levels, phloem loading and carbohydrate storage in various genotypes. Phloem apoplasmic and symplasmic pathways are shown, for our genotypes. Passive diffusion of sucrose occurs through plasmodesmata (PD) and water exchange across membranes are facilitated by aquaporins (AQP). In the COLLECT phloem, intercellular transport of sucrose is typically mediated by efflux from PPC (phloem parenchyma cells) *via* to SWEET facilitators, followed by influx in CCto (companion cells) or SE (sieve elements) *via* the SUC2 active sucrose/proton symporter, or into Mes (mesophyll cells) *via* SUC1. Sucrose can also be cleaved by invertases (INV) into hexoses ith influx by hexose transporters (HT). In the TRANSPORT phloem, sucrose that has leaked from the SE can be retrieved by SUC2 into the SE or stored in vascular cells (VS) after influx mediated by unidentified sucrose (SUC) or hexose transporters (HT). In this model, AQP are only indicated in cells where the apoplasmic water potential is higher than that of the cytosol and causes entry of water. *SUC2*, specifically expressed in the CC, is downregulated in response to high leaf sucrose levels, while *SUC1,* expressed in the mesophyll, is upregulated under the same conditions. **(a)** : In wild-type plants, loading results from efflux of sucrose into the apoplasm of the phloem parenchyma cells by SWEET 11 and SWEET12, its entry into the CC by SUC2, then passive diffusion through PD between CC and SE. Under normal conditions, SUC2 regulates the sucrose concentration in the apoplasm of the CC/SE complex, thereby maintaining a water potential difference across the plasma membrane of the CC/SE. This enables water uptake into the CC and the SE in the collection phloem, generating axial phloem mass flow. It also supports water uptake in the transport phloem in coordination with sugar retrieval along the transport pathway, limiting the storage of sugars in the VC. **(b)** : When SUC2 is inactive in the collection phloem, as in *miSUC* lines, increased sucrose concentration occurs in the apoplasm of the minor veins. This perturbation impacts the water potential and impairs water uptake in the SE/CC complex. The high levels of sucrose in the apoplasm, in turn, induce the upregulation of *SUC1*, increasing the sucrose influx in mesophyll cells and creating a gradient of sucrose concentration from the mesophyll to the phloem. This process shifts the phloem loading towards a symplasmic pathway, which maintains the phloem mass flow. **(c)** : When SUC2 is inactive in the transport phloem, as in *suc2pC* line, increased sucrose concentration occurs in the apoplasm of the CC and VC. This likely reduces water uptake in these cells, and carbohydrates in excess are preferentially stored as starch in the plastids. **(d)** : When *SUC2* is inactive in the collection and in the transport phloem, as observed in the *suc2* mutant, a dramatic increase in sucrose concentration occurs in the apoplasm of the phloem cells, which lowers apoplasmic water potentials and prevents the entry of water in phloem cells. The subsequent upregulation of *SUC1* in the mesophyll cells, increases sucrose influx in the mesophyll cells, and shifts the phloem loading towards a symplasmic pathway. However, the low osmotic potential in the apoplasm poses a challenge by restricting water entry across the plasma membranes of phloem cells. This limitation impacts phloem mass flow, both reducing phloem sap velocity and increasing sap viscosity. Carbohydrates in excess are stored as starch in the plastids. **(e)** : When SUC2 is active, the increased *NHL26* expression in the minor veins in the *GAS:NHL* line, impacts PD permeability, elevating sucrose levels in CC cytosol within the collection phloem. This reduces *SUC2* expression through a feedback mechanism, increasing sucrose concentrations in the apoplasm. However, the osmotic effect is insufficient to impede water entry in the minor veins, thereby maintaining the phloem mass flow. However, the high apoplasmic sucrose levels cause an upregulation of *SUC1*, increasing sucrose influx in the mesophyll cells and promoting an accumulation of soluble sugars in those cells, notably in the vacuole. **(f)** : When SUC2 is active, the overexpression of *NHL26* in *35S:NHL* line, alters plasmodesmata permeability, elevating sucrose levels in CC cytosol within the collection and transport phloem. This negatively affects *SUC2* expression and increases sucrose concentrations in the apoplasm. A low water potential in the apoplasm likely impedes water entry in the minor veins, resulting in reduced phloem mass flow. The excess sucrose accumulates in the phloem parenchyma cells and in mesophyll cells, where it is stored, as starch in the plastids and soluble sugars in the vacuole. Mes: Mesophyll cell, PPC: phloem parenchyma cell, CC: companion cell, SE: sieve element, VC: vascular cell. Phloem cells are in blue (CC, PPC and VC) and grey (SE), and mesophyll cells in green (Mes). The intensity of the color (blue, green or grey) represents the levels of sucrose in the cell, with a light color corresponding to a low level and an intense color to a high level. SUC: sucrose transporter, HT= hexose transporter, SWT: SWEET11/12 facilitator, AQP=aquaporin, cwINV= cell wall invertase, cINV=cytosolic invertase, vINV=vacuolar invertase. Open plasmodesmata are represented in blue, closed plasmodesmata occluded by NHL26 in black. ψ_apo_: water potential of the apoplasm.

The SUC2 closest ortholog, SUC1, has a similar affinity for sucrose (Sauer & Stolz, 1994) and complements *suc2* mutant for sucrose influx (Wippel & Sauer, 2012). The function of SUC1 in source organs remains unexplored, and there are conflicting reports regarding its expression in leaves. Recent leaf single-cell transcriptomics and translatome studies indicate that *SUC1* is expressed in the mesophyll cells (Mustroph *et al*., 2009; Kim *et al*., 2021; Xu & Liesche, 2021). In contrast to *SUC2*, which is downregulated by sucrose (Solfanelli *et al*., 2006), *SUC1* is upregulated (Solfanelli *et al*., 2006). Consistently, the high sucrose levels in the leaves of *miSUC* and *suc2pC* plants were associated with an increased expression of *SUC1* compared to the wild-type. At the same time, the phloem exudation rate was maintained. Our data suggest that *SUC1* likely functions in the mesophyll cells to retrieve excess sucrose from the apoplasm, similar to its proposed role in the roots (Durand *et al*., 2018).

In the two *miSUC* lines, defective for *SUC2* expression in the minor veins, the retrieval of sucrose by SUC1 may increase cytosolic soluble sugar concentrations in the cells at the periphery of the minor veins while the starch levels remain stable. The downregulation of *HXK1* and *HXK2* and upregulation of *cINV1/2* in *miSUC plants* confirm high sucrose levels in the cytosol. Because phloem transport is maintained in these lines, we propose that the accumulation of sugars may establish a sucrose concentration gradient from the mesophyll to the vascular cells (Fig. **8c**), thereby enhancing diffusion via bulk flow through PD along this gradient, a mechanism proposed for symplasmic loaders (Schulz, 2015). Interestingly, in the *suc2pC* line, which lacks *SUC2* expression in the transport phloem – i.e. the main veins - we also observed an upregulation of *SUC1,* with high starch accumulation in the leaf, and upregulation of G6PT2 and APL3, indicating that in this case, high sucrose in the apoplasm also leads to an increased storage capacity for starch. This suggests that the levels of apoplasmic sugars in the transport phloem may also promote starch storage in the plastids.

These findings support Turgeon’s hypothesis (Turgeon, 2010), that active loading evolved not primarily to facilitate phloem transport but to enable plants to utilize foliar carbon reserves. Consequently, both *miSUC* and *suc2pC* lines, despite maintaining a normal phloem sugar exudation rate, accumulated higher levels of non-structural carbohydrates. This accumulation was associated with slower growth rates and reduced rosette and stem growth, consistent with the growth-storage trade-off paradigm (Martínez-Vilalta *et al*., 2016).

### Apoplasmic sucrose levels and water flows in the phloem

The *suc2* mutant also showed high starch accumulation, reduced rosette and stem growth, high sugar content in leaf AWF, and high *SUC1* expression compared to the wild-type. The data support the hypothesis that excess sucrose in the apoplasm, uptaken by SUC1 in mesophyll cells, creates a cytosolic sucrose concentration gradient, from the mesophyll to the vascular cells (Fig. **8d**). We propose that such a gradient facilitates bulk flow diffusion of sucrose through PD, driving symplasmic loading. This hypothesis is supported by the observation that the *suc1suc2* double mutant exhibits an additive phenotype in plant growth (Fig. **S5**), indicating that sucrose uptake by SUC1 in the mesophyll cells becomes essential in the absence of SUC2 activity, while maintaining phloem loading to some extent.

Most intriguingly, we observed high sugar amounts in the exudates of the *suc2* plants, revealing dramatic consequences of the loss of *SUC2* expression on phloem transport activity by contrast to the *SUC2* downregulated *miSUC* and *suc2pC* plants. It is reasonable to assume that this is due to a high sugar concentration in the phloem sap. If so, changes in sucrose concentration could contribute to increase phloem sap viscosity and reduce phloem sap flow rate, which has been confirmed experimentally (Gottwald *et al*., 2000), with a doubling in transit time between organs in *suc2*. This hypothesis is also consistent with earlier ^14^C labeling experiments with *suc2*, which demonstrated reduced carbon allocation to the roots (Gottwald *et al*., 2000) and reduced ^14^C level in the exudates (Srivastava *et al*., 2008). The increase in sucrose concentration might be caused, in part, by the combination of high sugar content in the mesophyll cells, revealed by the high carbohydrate storage in the leaf - and subsequent high symplasmic bulk flow from mesophyll to phloem cells.

However, the high sucrose concentration in the suc2 mutant might also result from the reduced entry of water by osmosis in the phloem cells, because of high sucrose concentration in the apoplasm, reducing dramatically phloem mass flow, which is driven by an osmotically generated pressure gradient between the apoplasm and the cytosol. We showed a high sugar concentration in the AWF of *suc2*, which may minimize the osmotic potential difference across the plasma membrane of the PPC and CC/SE complex, thereby limiting water uptake. A reduced radial water flow potentially leads to elevated sucrose concentration in the sap and increased viscosity, reducing phloem flow rate. We propose that a combination of low flow velocity and high sap viscosity increases the turgor pressure of the SE in *suc2* (Fig. **8d**), explaining the very high rates of sucrose exudation in *suc2* after sectioning the highly pressurized sieve tubes.

Our findings provide new insights into the *suc2* mutant phenotype. Gould et al. (Gould *et al*., 2012) suggested that the absence of SUC2 in the collecting phloem reduces turgor pressure in SE and slowing down transport. However, the elevated sugar exudate rates and sugar levels in the apoplasmic fluids of *suc2* contradict Gould’s hypotheses, suggesting high osmotic potentials in both compartments. Contrary to Gould’s proposals, our findings imply that *SUC2 loss of function*-mediated elevation of apoplasmic sucrose content in the phloem impedes the water flow that drives phloem flow.

### Sap viscosity, turgor pressures and phloem unloading

Another feature of *suc2* plants was a reduction in seed weight and lower C, protein and lipid contents in the seeds, effects that were also observed in *suc2pC* plants, compared to the wild-type, which suggests an impairment in phloem unloading in sink organs. Interestingly, as observed in young seedlings *in vitro* (Gottwald *et al*., 2000), *suc2* root growth is severely reduced compared to wild-type, which is partially mitigated in a sugar-rich environment (Fig. **S6**). Unloading of solutes in the root is mainly symplasmic, through funnel-PD at the root tip, driven by a combination of mass-flow and diffusion through PD (Ross-Elliott *et al*., 2017). Our findings of high sugar in the AWF and high rate of sugar exudation suggest high sap viscosity in the sieve tubes and a high turgor pressure. Interestingly, as observed in young seedlings *in vitro* (Gottwald *et al*., 2000), root growth is partially mitigated in a sugar-rich environment (Fig. **S6**). Several studies have shown that the PD permeability is osmo-regulated in response to turgor pressure exerted on both sides of the PD (Hernández-Hernández *et al*., 2020). When osmolyte concentrations, such as sucrose, become excessively high on one side, plasmodesmata close, disrupting symplasmic connectivity. Our findings indicated that in a hypertonic sugar-rich environment, there may be less disparity in the osmotic potential of the root outer cell layers and in phloem cells. Consequently, a hypertonic environment should restore PD connectivity and facilitate symplasmic unloading at the root tip, potentially accounting for the observed increase in root growth in a sugar-rich environment.

### Impacts of apoplasmic and symplasmic loading strategies on foliar carbon storage

The significant increase in sucrose content observed in the AWF of the *NHL* and *SUC* lines compared to the wild-type was associated with an increased expression of *cwINV1 and SUC1*, two genes expressed in mesophyll cells (Kim *et al*., 2021; Xu & Liesche, 2021) and with elevated foliar soluble sugars. These results suggest that even minor reductions in symplasmic or apoplasmic pathways increase foliar carbon storage in the mesophyll. Correspondingly, genes involved in photosynthesis (e.g., *RBCS* and *LHCB1*) or cytosolic glucose signaling (e.g., *HXK1* and *HXK2*) exhibited slight downregulation in some lines, aligning with negative feedback regulation of photosynthesis by high sugar levels (Griffiths *et al*., 2016). The data highlight shared metabolic and photosynthetic responses to impaired *SUC2-* dependent apoplasmic or impaired symplasmic sugar loading. Additional genes like *FRK2*, *FRK6*, *SWEET11,* and *SWEET12,* expressed in phloem cells, are consistently upregulated across all lines, suggesting a subsequent impact on the pools of cytosolic soluble sugar in vascular tissues.

We observed marked differences in the foliar carbon storage of the plants impaired in the symplasmic pathway, *35S:NHL* and *GAS:NHL,* with a moderate soluble sugar accumulation compared to the plants impaired in the SUC2-apoplasmic pathway. Remarkably, it was also associated with a reduction in the sucrose exudation rate in *35S:NHL* plants, with a similar trend observed in *GAS:NHL plants*, in contrast to no alteration in exudate rates in *miSUC* and *suc2pC* plants. This finding rules out the hypothesis that the reduction was due to impaired *SUC2*-mediated apoplasmic phloem loading, which maintained the sucrose exudation rate. Instead, we propose that PD-dependent sucrose transport between the CC and the SE is also disrupted in the *GAS:NHL* line (Fig. **8e,f**), thus reducing phloem loading.

Interestingly, the foliar accumulation of soluble sugars in *GAS:NHL* and *35S:NHL* plants correlates with the up-regulation of *vINV1* and/or *vINV2*, key players in vacuolar sucrose turnover required for proper plant development (Vu *et al*., 2020). This suggests that the symplasmic pathway influences sucrose homeostasis within the vacuole. In contrast, in lines with impaired *SUC2* expression the observation of a downregulation of *vINV1*, *vINV2*, and *SWEET17* implies distinct consequences for sugar partitioning when apoplasmic pathway is impaired. However, the active tonoplastic sugar transporter genes *TMT1/2* remained unaffected, indicating a complex interplay in sugar partitioning between the apoplasm and the vacuole — an aspect that has been inadequately explored thus far. Based on these observations, we propose that the symplasmic and *SUC2*-dependent apoplasmic pathways differentially impact foliar carbon storage, favoring either sugar storage in the vacuole or starch in the chloroplasts, depending on apoplasmic sugar levels and water flows.

### Conclusions

Our findings confirm the pivotal role of *SUC2* in regulating phloem loading and unloading. The data support the hypothesis that SUC2 achieves this by affecting sucrose levels in the apoplasm, thereby influencing water potential gradients and impacting water flow into or out of the phloem cells. Moreover, the data suggest that differences in osmotic potential between the apoplasm and the cytosol, responsible for water entry in the CC/SE complex, may alter phloem loading. This, in turn, may affect the permeability of PD for unloading, as suggested by Yan & Liu (Yan & Liu, 2020).

The suggested collaborative role of *SUC2*, working alongside *SUC1* to regulate sugar content in the apoplasm and to control water flow to the phloem, could have significant implications for leaf water status and photosynthesis. It has long been recognized that sugar levels in the apoplasm, including sucrose and glucose, influence guard cell regulation, stomatal opening, and gas exchanges (Daloso *et al*., 2016; Flütsch & Santelia, 2021). Whether the activity of SUC1/SUC2 in apoplasmic sugar regulation contributes to the negative feedback regulation of sugars on photosynthesis remains a subject for future investigation.

## Acknowledgments

We thank Hervé Ferry and Joël Talbotec for taking care of the plants in the greenhouse facility. We thank Roua Jeridi for her help to the initial collects of apoplasmic washing fluids. We thank the Plant Observatory. This work has benefited from the support of IJPB’s Plant Observatory technological platforms. The IJPB benefits from the support of Saclay Plant Sciences-SPS (ANR-17-EUR-0007). We thank Dr Michael Thorpe for his helpful comments that improved the manuscript. We thank Bryan Ayre and Nathalie Pourtau for the gift of seeds and plasmids.

## Author Contributions

F.V. and S.D. conceived and supervised the experiments. F.V. and L.B. realized the experiments. First draft was done by S.D. Editing was done by S.D., F.V., R.L.H. and C.B.

## Data availability

Raw data and seeds from Arabidopsis lines supporting the findings of this study are available from the corresponding author SD on request.

## Supporting Information

**Fig. S1.**
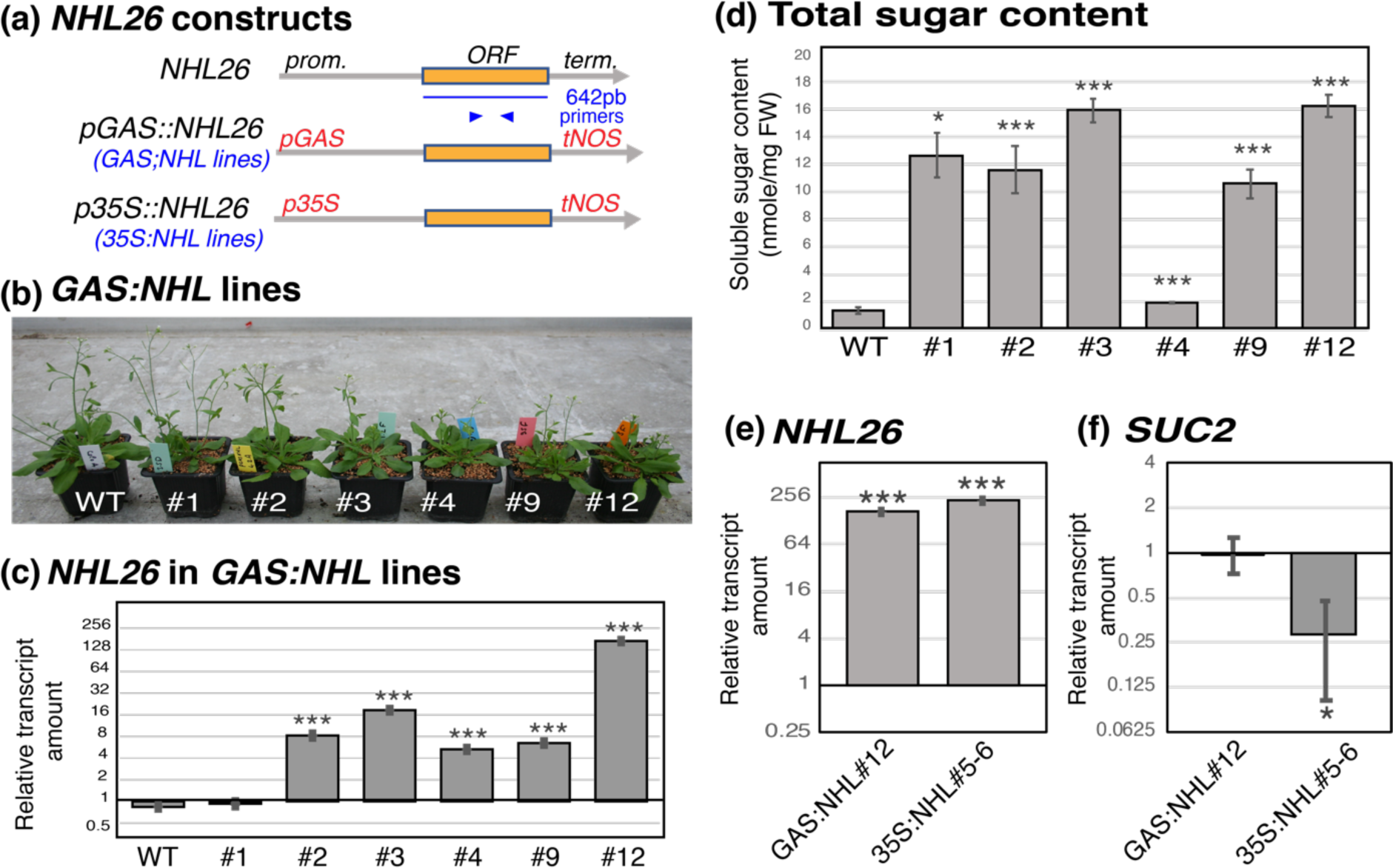
Characterization of the *NHL*-lines. **(a)** Schematic representation of *NHL26* constructs. On the top panel: schematic representation of the structure of the *NHL26* gene (AGI number At5g53730). The length of the exon and the position of the primers used for RT-QPCR are indicated below. prom.: promoter, term.: terminator. On the middle panel, representation of the *pGAS::NHL26* construct. The coding region of *NHL26* was fused to the promoter of the *Cucumis melo* galactinol synthase gene to drive gene expression specifically in the minor veins of the mature leaves, as described (Srivastava *et al*., 2008). On the bottom panel, representation of the *p35S::NHL26* construct (Vilaine *et al*., 2013). The coding region of *NHL26* was fused to the cauliflower mosaic virus (CaMV) 35S constitutive promoter. **(b)** to (**f**) Phenotypes of 5 weeks-old plants, grown in long-day condition: **(b)** Phenotype of representative plants from 6 independent *pGAS::NHL26* transgenic lines (hereafter referred to as *GAS:NHL*). **(c)** Relative *NHL26* transcript amount in the *GAS:NHL* transgenic lines. The data are shown on a log_2_ scale after normalization by the accumulation of *NHL26* transcripts in WT plants. **(d)** Accumulation of total soluble sugars in the *GAS:NHL* transgenic lines. The total sugar contents (glucose, fructose plus sucrose) are expressed in nmoles per mg of fresh weight (FW). **(e)** and **(f)** Relative *NHL26* and *SUC2* transcript amounts in the *GAS:NHL*#12 and *p35S::NHL26#5-6* (hereafter referred to as *35S:NHL*) plants. The transcript amount was assessed by qRT-PCR, normalized relative to that of the reference gene *TIP41*, and shown as fold-change compared to wild-type plants, on a log_2_ scale. Bar plots and error bars represent the mean and *se* (*n* = 6). Asterisks indicate significant differences compared to control plants (* p < 0.05; ** p < 0.01; *** p < 0.001)

**Fig. S2.**
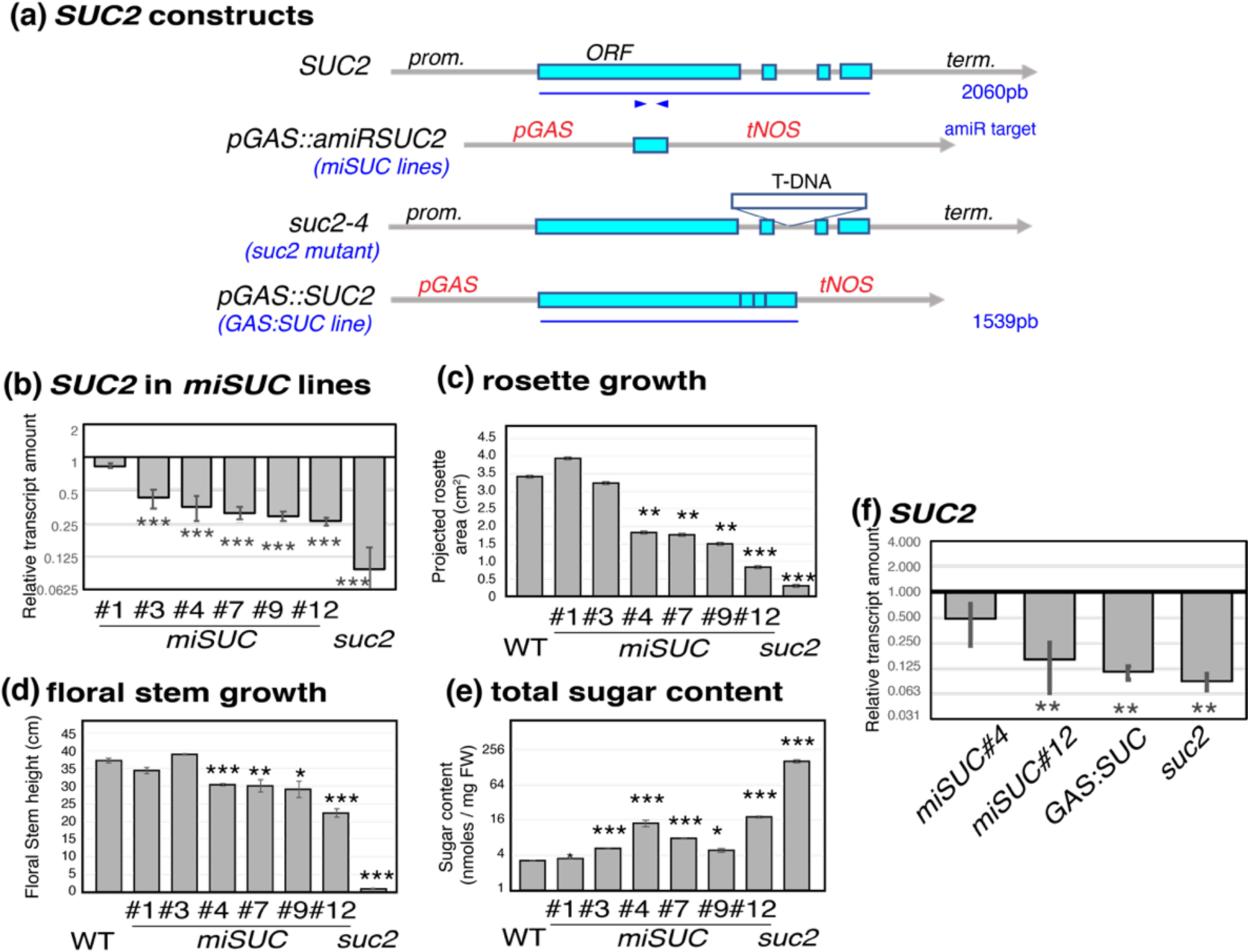
Characterization of the *SUC*-lines. **(a)** Schematic representation of *SUC* constructs. On the top panel: schematic representation of the structure of the *SUC2* gene (AGI At1g22710). The length of the coding region (including exons and introns) and the position of the sequence targeted by the *amiRNA* are indicated below. prom.: promoter, term.: terminator. Below, representation of the *pGAS*::*amiR:SUC2* construct. A microRNA targeting *SUC2* was fused to the promoter of *CmGAS1* to drive microRNA expression specifically in the minor veins of the mature leaves. On the middle panel, representation of the *suc2-4* mutant. On the bottom panel, representation of the *pGAS::SUC2* construct used to partially restore *SUC2* gene in the minor veins of the *suc2-4* mutant. **(b)** to (g) phenotypes of plants grown in long-day conditions **(b)** Relative *SUC2* transcript amount in representative plants from 6 independent *amiR::SUC2* (hereafter referred to *miSUC)* transgenic lines. The transcript amount was assessed by qRT-PCR, normalized relative to that of the reference gene *TIP41*, and shown as fold-change compared to wild-type plants, on a log_2_ scale. **(c)** Rosette growth of the representative plants from 6 independent *miSUC* transgenic lines. The projected rosette area was measured at 3 weeks on plants grown in a growth chamber in long-day condition. **(d)** Floral stem growth of the representative plants from 6 independent *miSUC* transgenic lines. The height of the floral stem was measured at 5 weeks on plants grown in a growth chamber in long-day condition. **(e)** Accumulation of total soluble sugars (glucose, fructose plus sucrose) in representative plants from 6 independent *miSUC* transgenic lines. Sugar contents are expressed in nmoles per mg of fresh weight (FW), shown on a log_2_ scale. **(f)** Relative *SUC2* transcript amounts in the *miSUC#4, miSUC#12, suc2pC and suc2* plants. The transcript amount was assessed by qRT-PCR, normalized relative to that of the reference gene *TIP41*, and shown as fold-change compared to wild-type plants, on a log_2_ scale. Bar plots and error bars represent the mean and *se* (*n* = 6). Asterisks indicate significant differences compared to control plants (* p < 0.05; ** p < 0.01; *** p < 0.001).

**Fig. S3.**
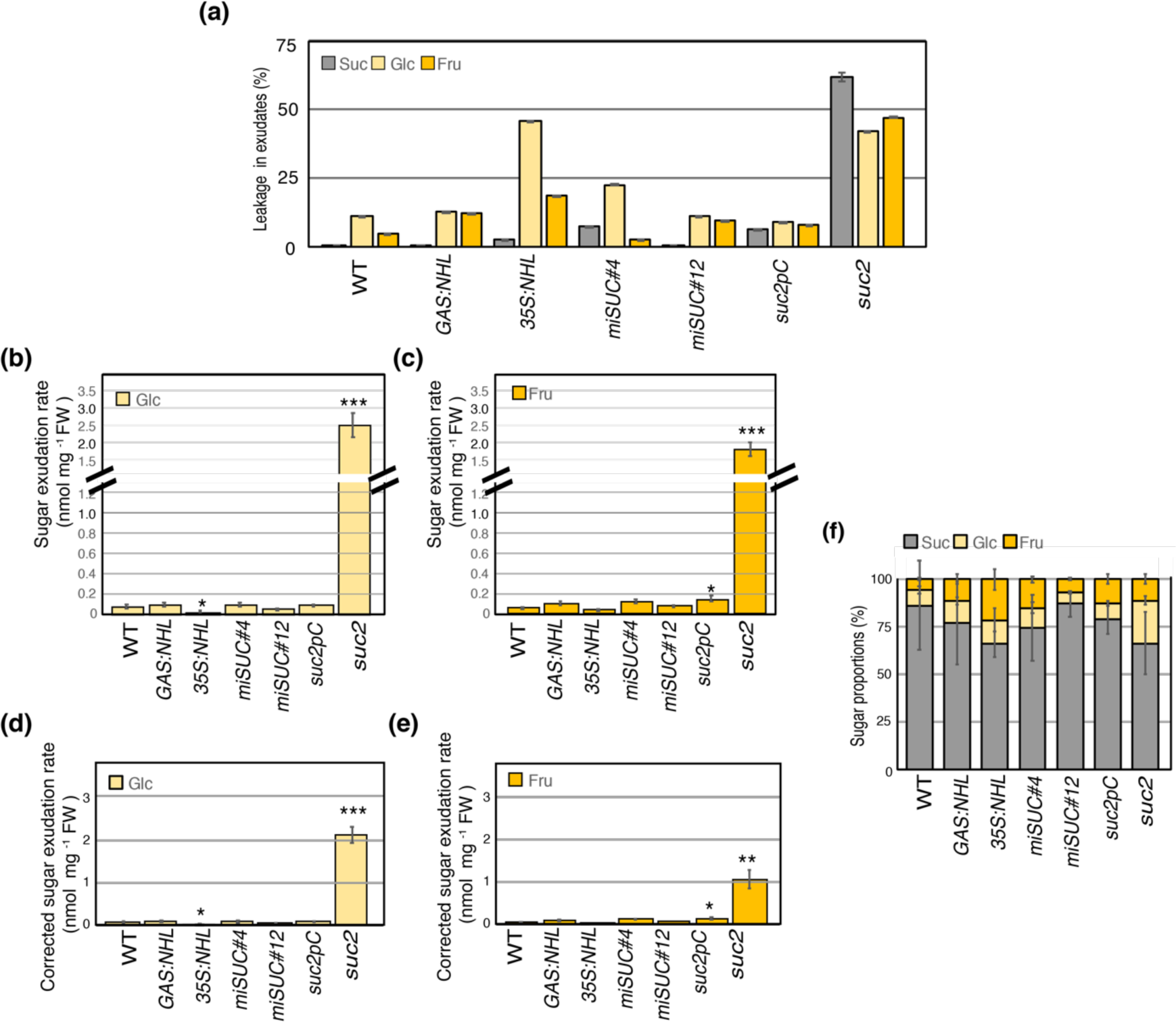
Rate of phloem hexoses and amino acids exudation in *NHL*- and *SUC*-lines. **(a):** Sugar leakage was measured on the exudate on a second rosette leaf of the same plant (5^th^ or 6^th^ leaf) collected using the same method but omitting EDTA, for 2 hours of exudation. **(b)** and **(c):** Glucose and Fructose exudation rate was measured on the phloem exudate collected by EDTA-facilitated exudation of one rosette leaf per plant (5^th^ or 6^th^ leaf) for 2 hours of exudation. **(d)** and **(e):** Corrected Glucose and Fructose exudation rates. **(f):** Proportion of disaccharide (sucrose) and monosaccharide (glucose and fructose) in the phloem exudates, after correction for leakage. Bar plots and error bars represent the mean and *se* (*n* = 6). Asterisks indicate significant differences compared to control plants (* p < 0.05; ** p < 0.01; *** p < 0.001). FW: fresh weight.

**Fig. S4.**
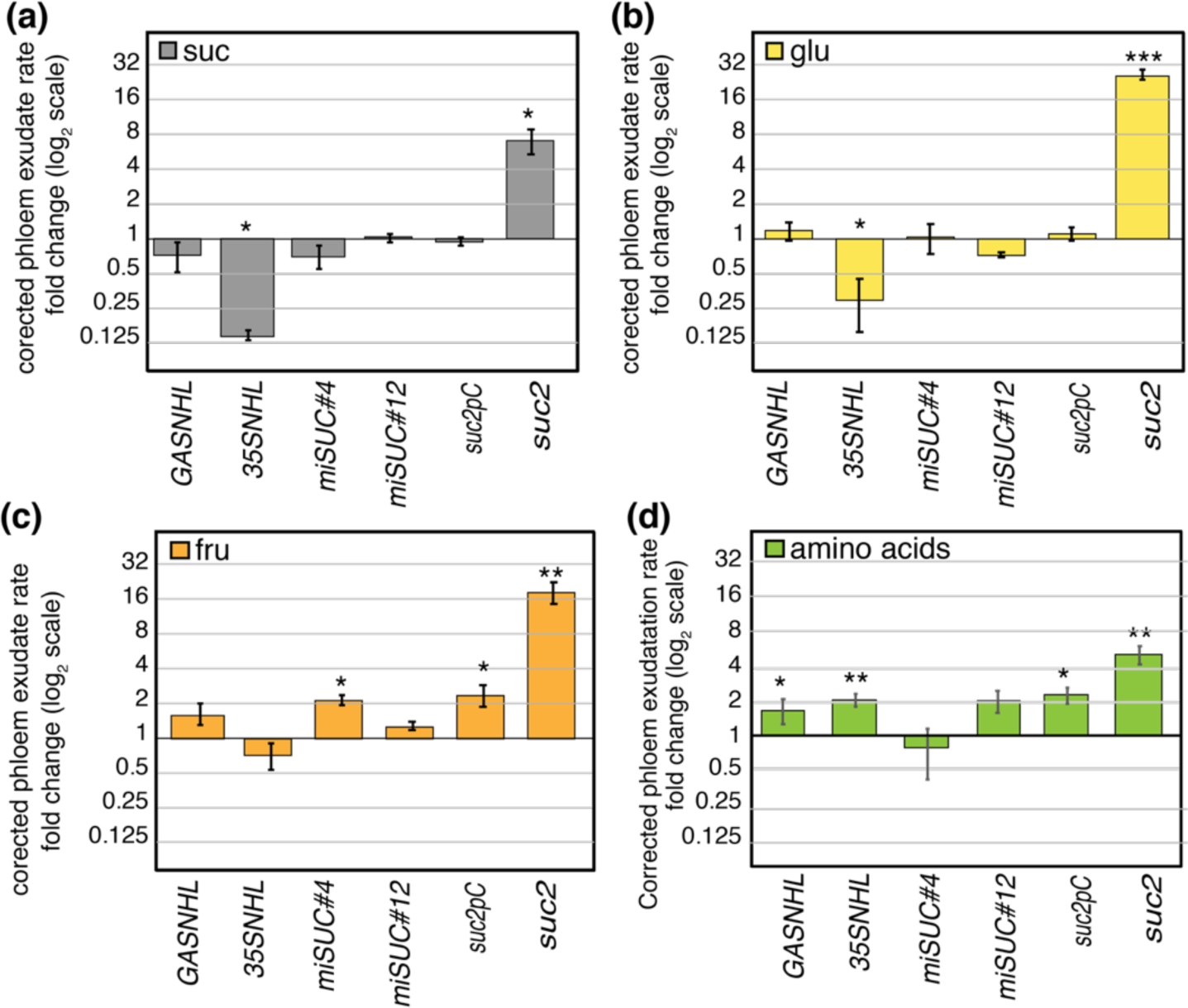
Fold-changes compared to WT in the exudation rates for sugars and amino acids in *NHL*- and *SUC*-lines. Fold changes compared to WT in the corrected exudation rate of **(a)** Sucrose, **(b)** Glucose, **(c)**, Fructose and **(d)** Total amino acids sugars, calculated by subtracting sugar leakage from the exudation rate of one rosette leaf per plant (5^th^ or 6^th^ leaf). Bar plots and error bars represent the mean and *se* (*n* = 6). Asterisks indicate significant differences compared to control plants (* p < 0.05; ** p < 0.01; *** p < 0.001).

**Fig. S5.**
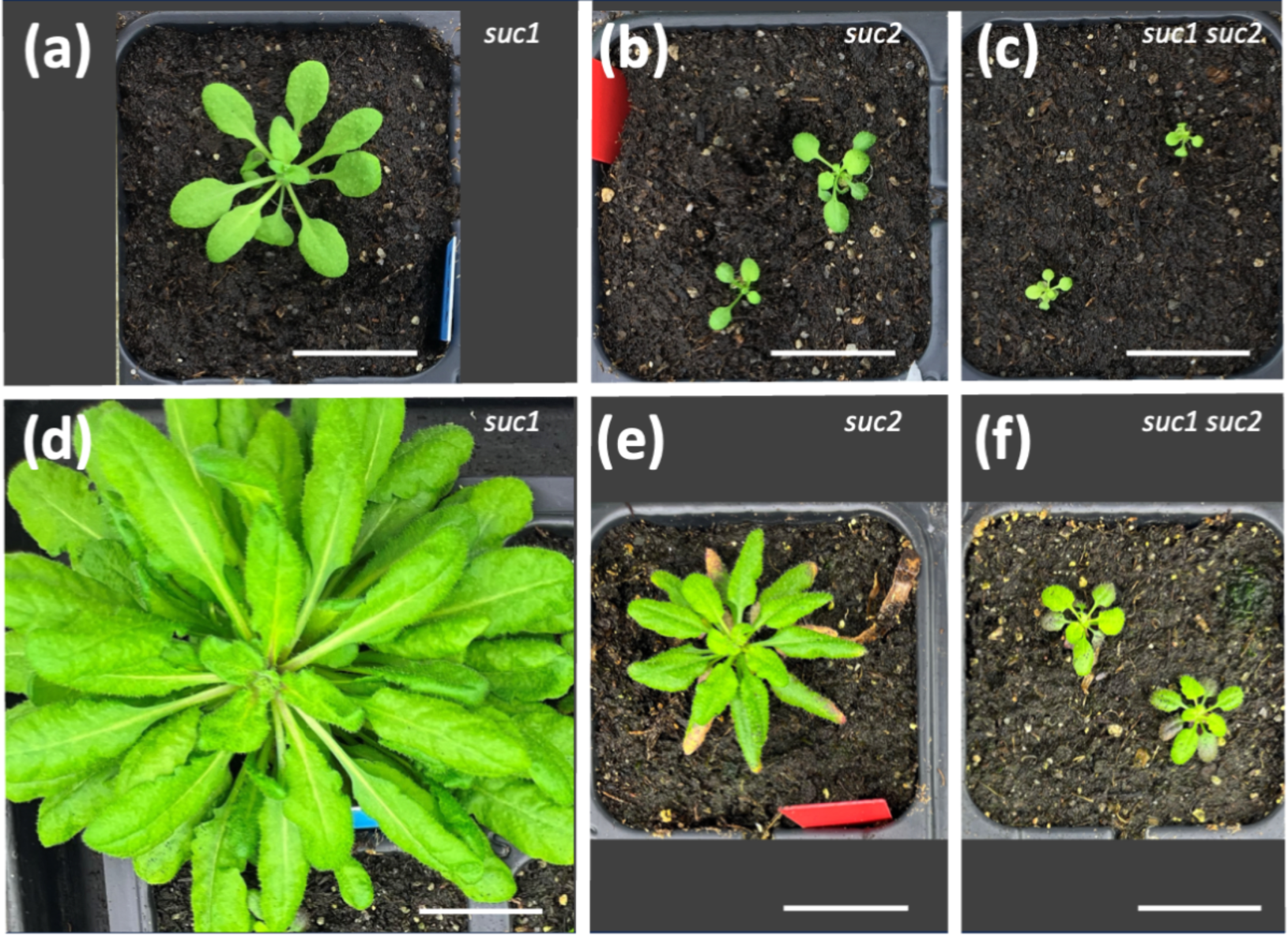
Phenotype of the *suc1* and *suc2* simple and double mutants. **(a)** to **(c)** 8 weeks-old plants; (**d**) to (**f**) 12 weeks-old plants, with *suc1* in (**a**) and (**d**), *suc2* plants in (**b**) and (**e**), and *suc1 suc2* plants on (**c**) and (**f**).Plants were grown under short days, to take into account that the plant phenotype tends to be milder for *suc2* when grown under shorter days, as already reported (Srivastava *et al*., 2009). Bar: 2 cm.

**Fig. S6.**
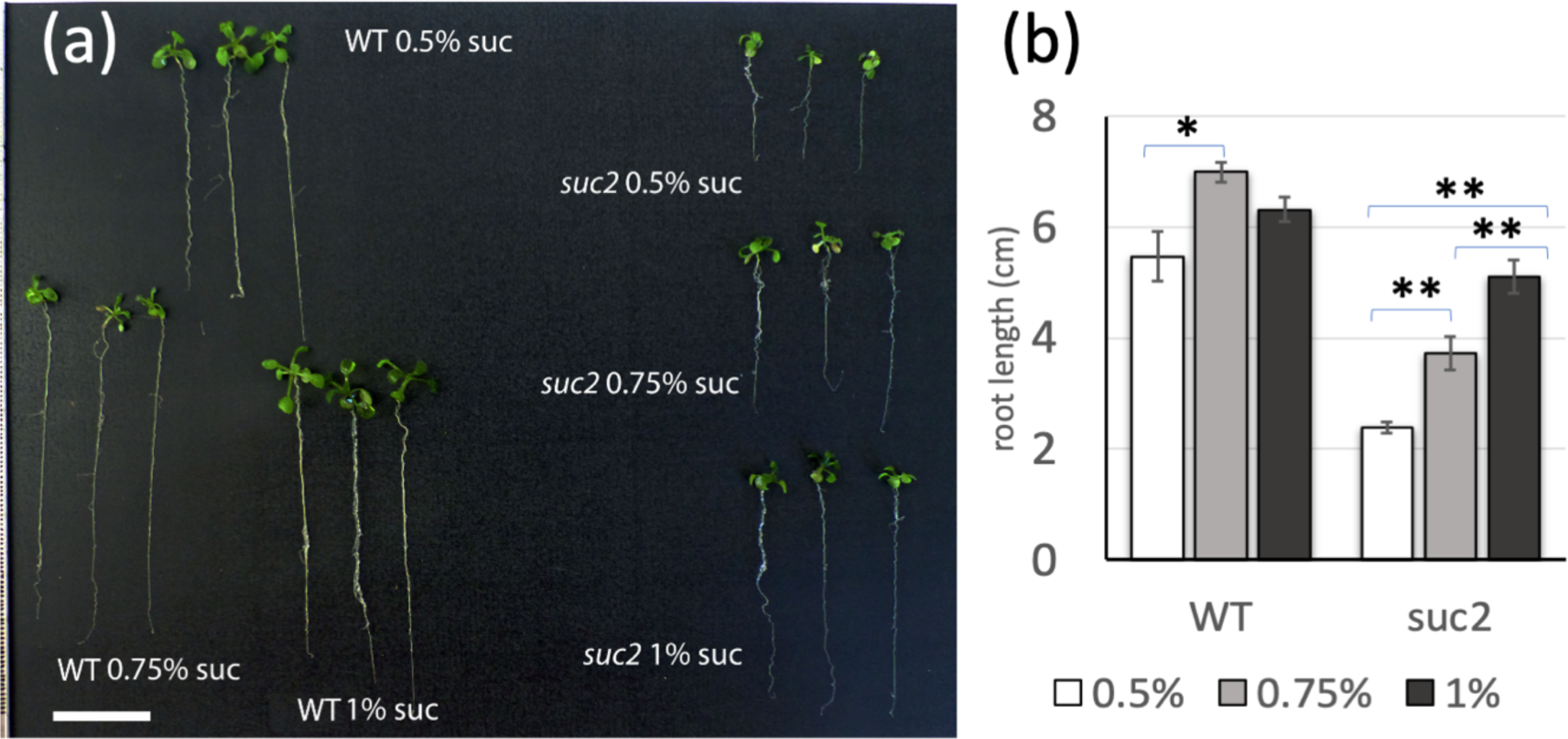
*In vitro* root growth of WT and *suc2* seedlings. Root growth of WT and *suc2* seedlings grown *in vitro*. **(a)** Plantlets were grown for 15 days on growth medium supplemented with 0.5%, 0.75% or 1% of sucrose (bar = 2 cm). **(b)** Root length (*n*=3 bar: mean +/- *SD*).

**Table S1.**
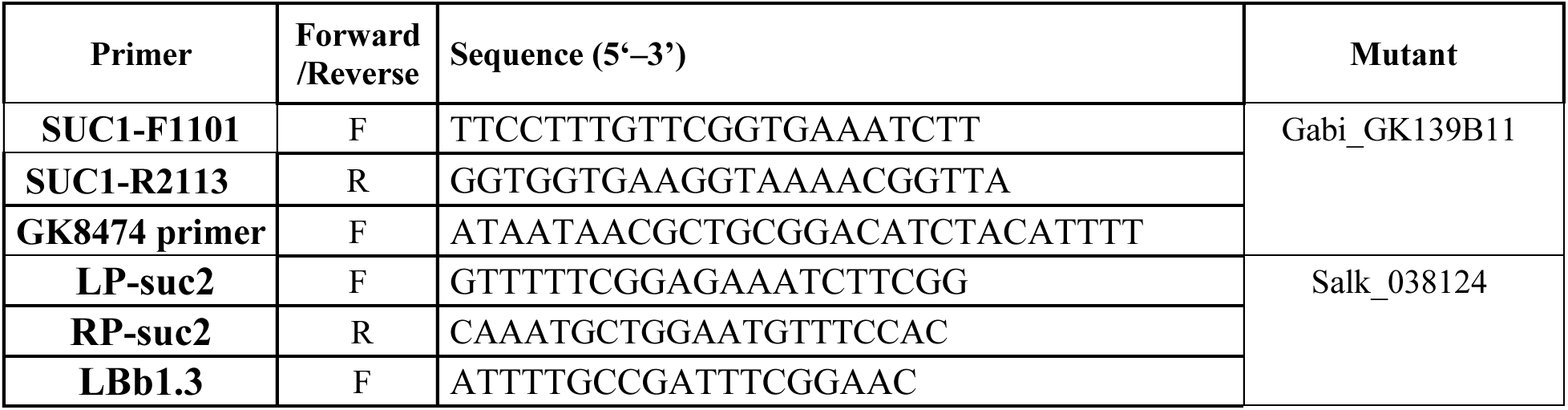
Primers for genotyping.

**Table S2.**
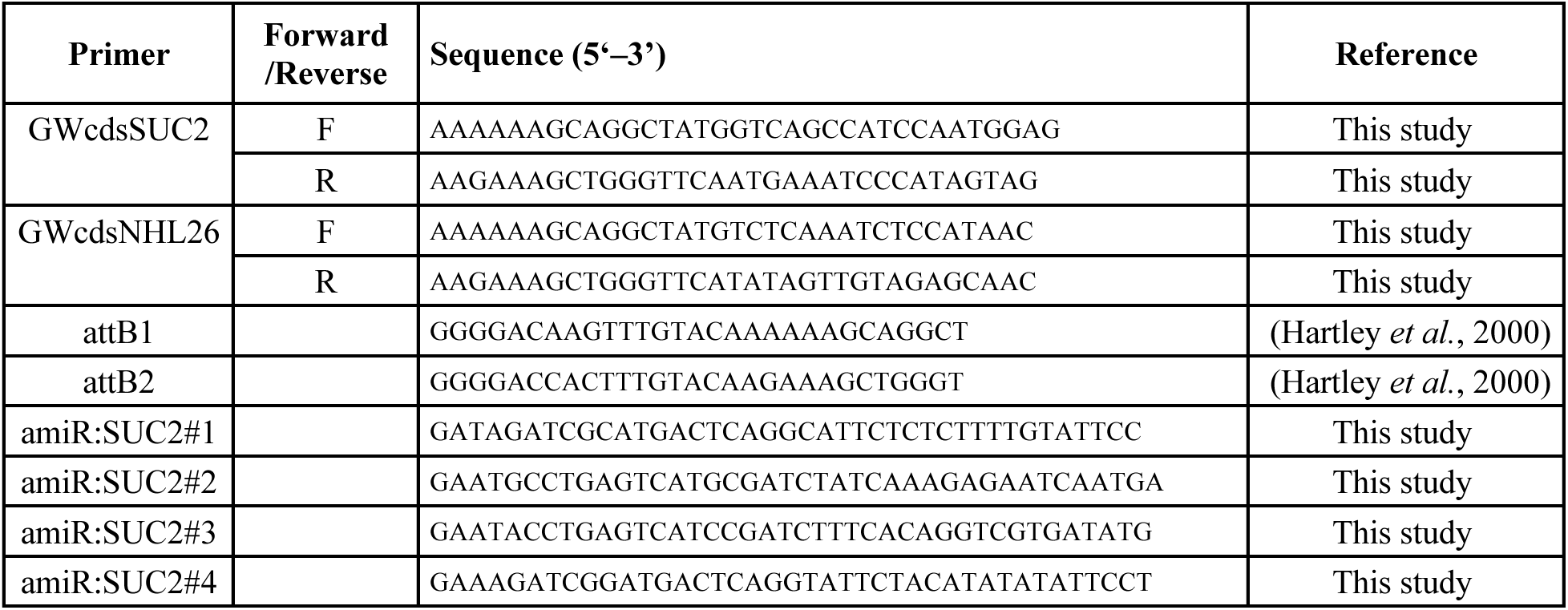
Primers for Gateway^R^ cloning.

**Table S3.** Vectors for cloning.

| Plasmids |  | Reference |
| --- | --- | --- |
| **pIPK-pGAS-R1R2-tNOS** | Gateway binary destination vector, with Galactinol synthase promoter and NOS terminator | This study |
| **pGWB2** | Gateway binary destination vector, with CaMV 35S promoter and NOS terminator | (Nakagawa *et al.*, 2007) |
| **pRS300** | Backbone for expressing plant artificial miRNAs | (Schwab *et al.*, 2006) |
| **pSG3K101** | Galactinol synthase promoter | (Haritatos *et al.*, 2000) |
| **pIPKb001** | Gateway destination vector | (Himmelbach *et al.*, 2007) |

**Table S4.**
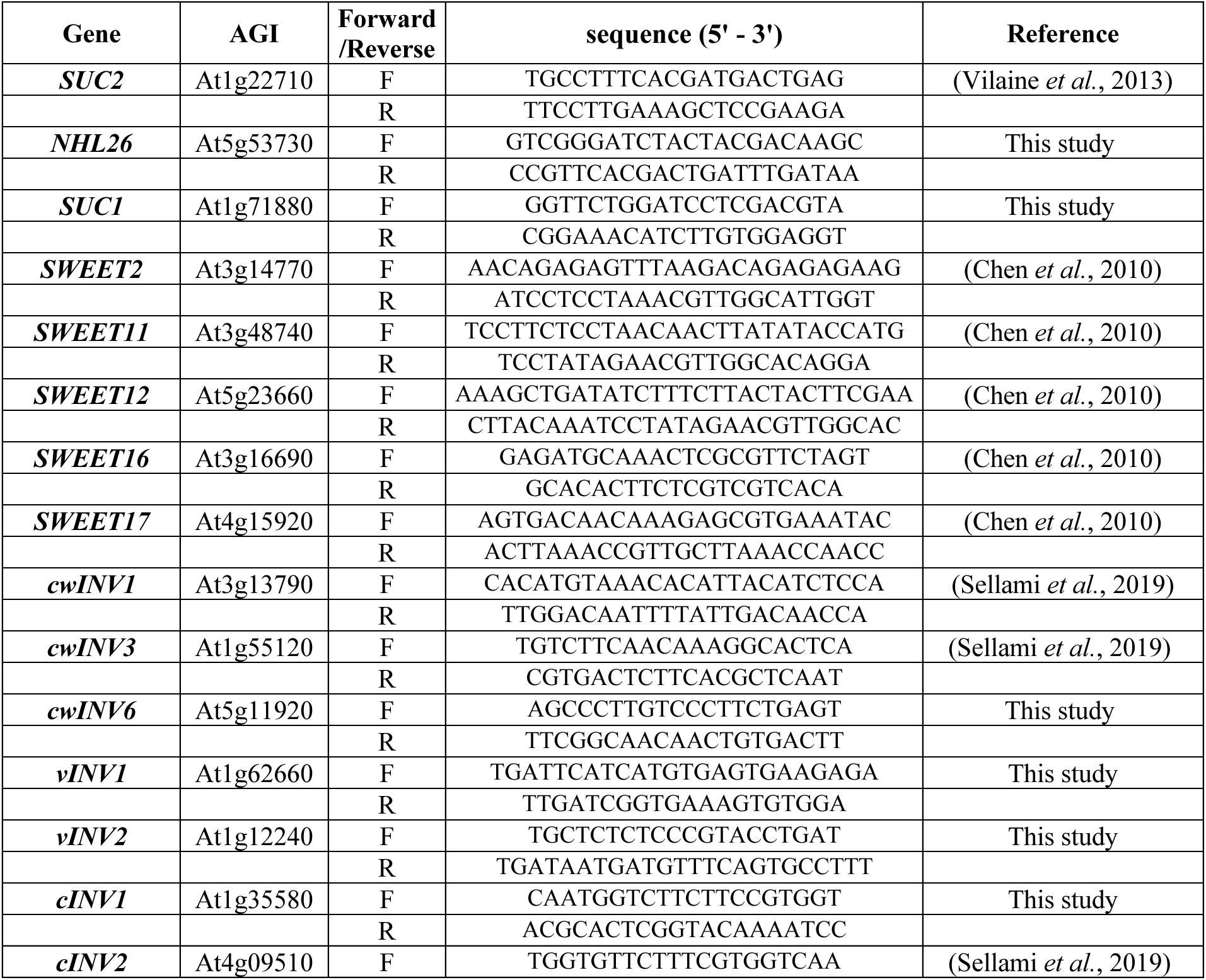
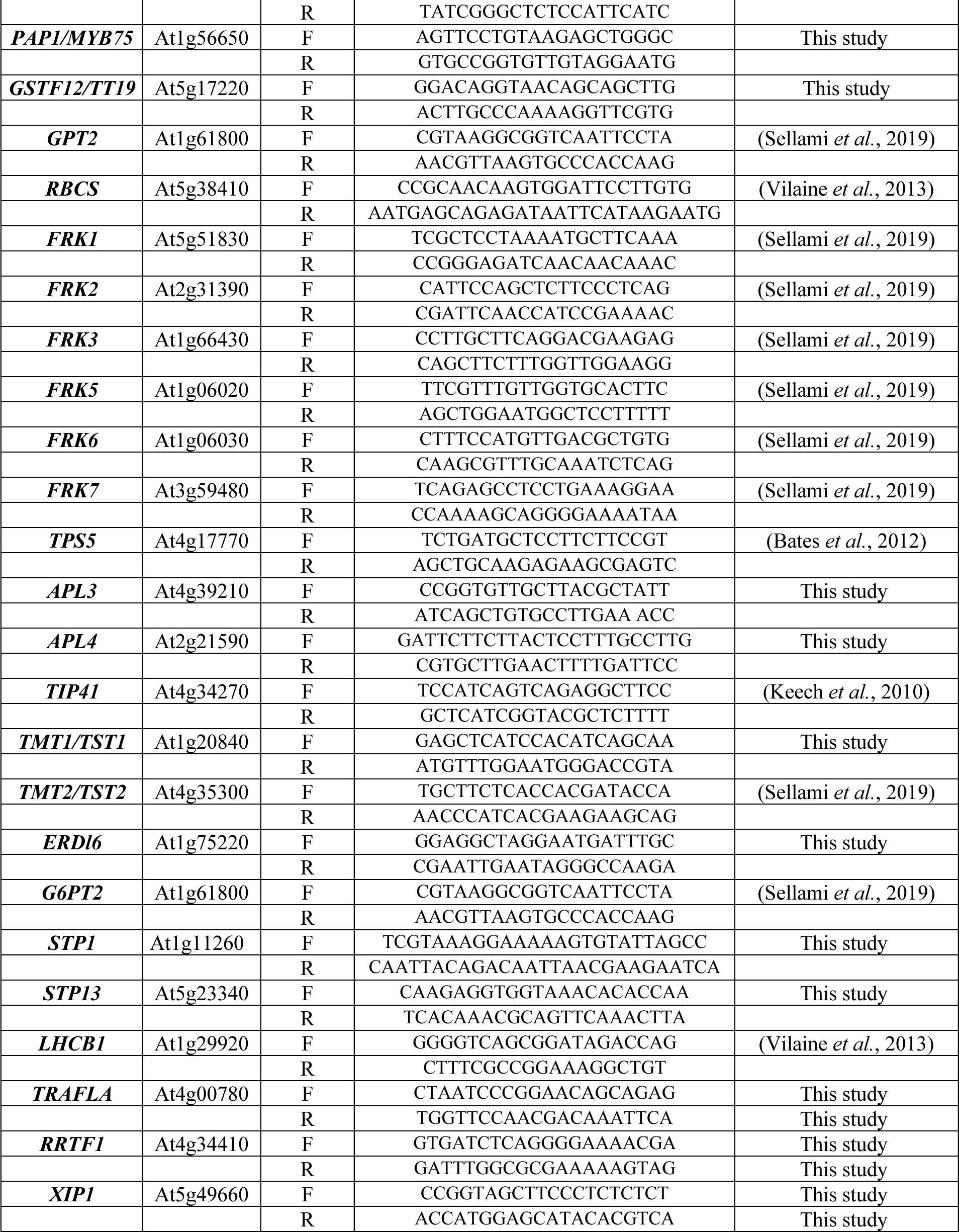
Primers for quantifying genes by RT-qPCR.

**Table S5.**
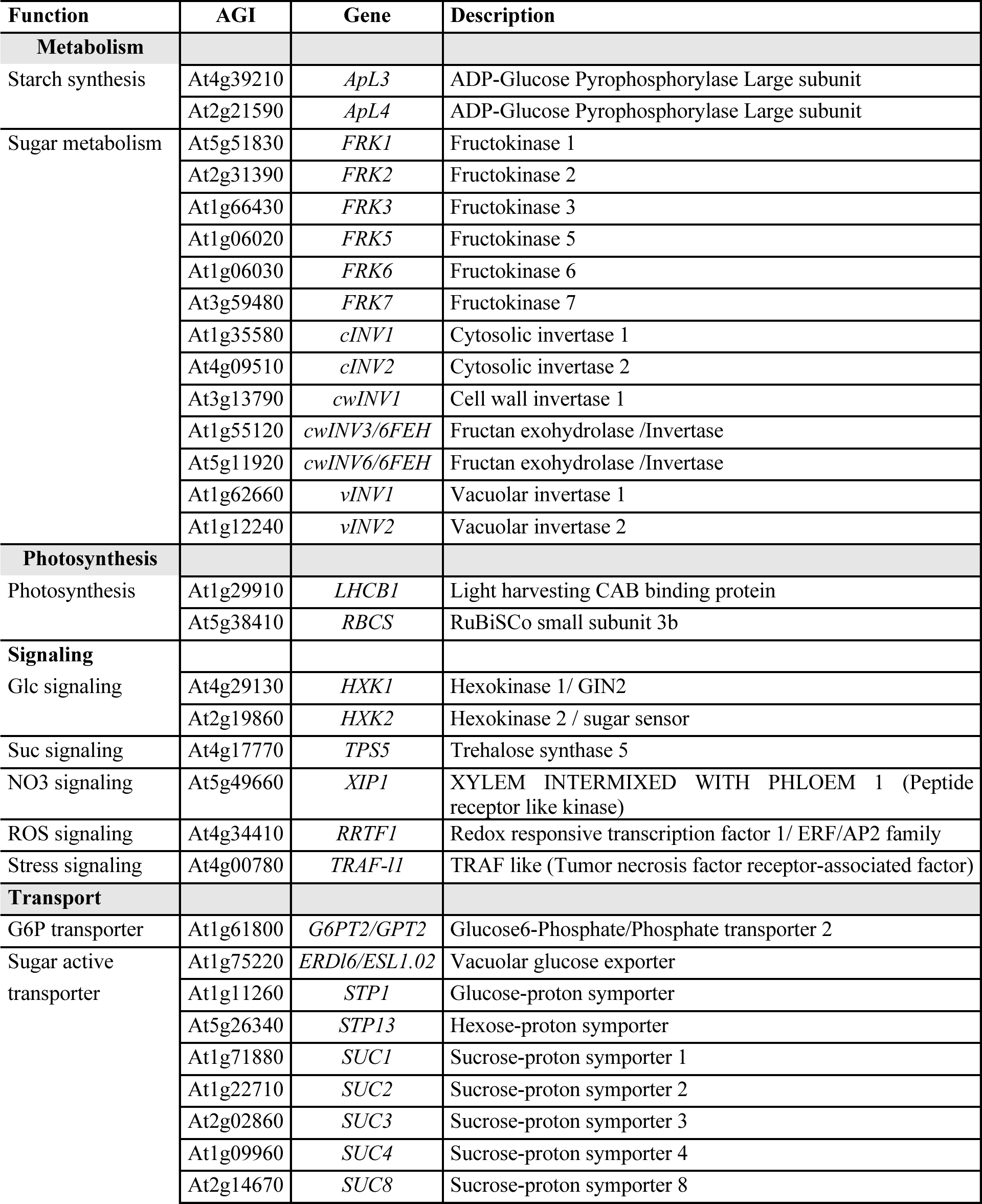
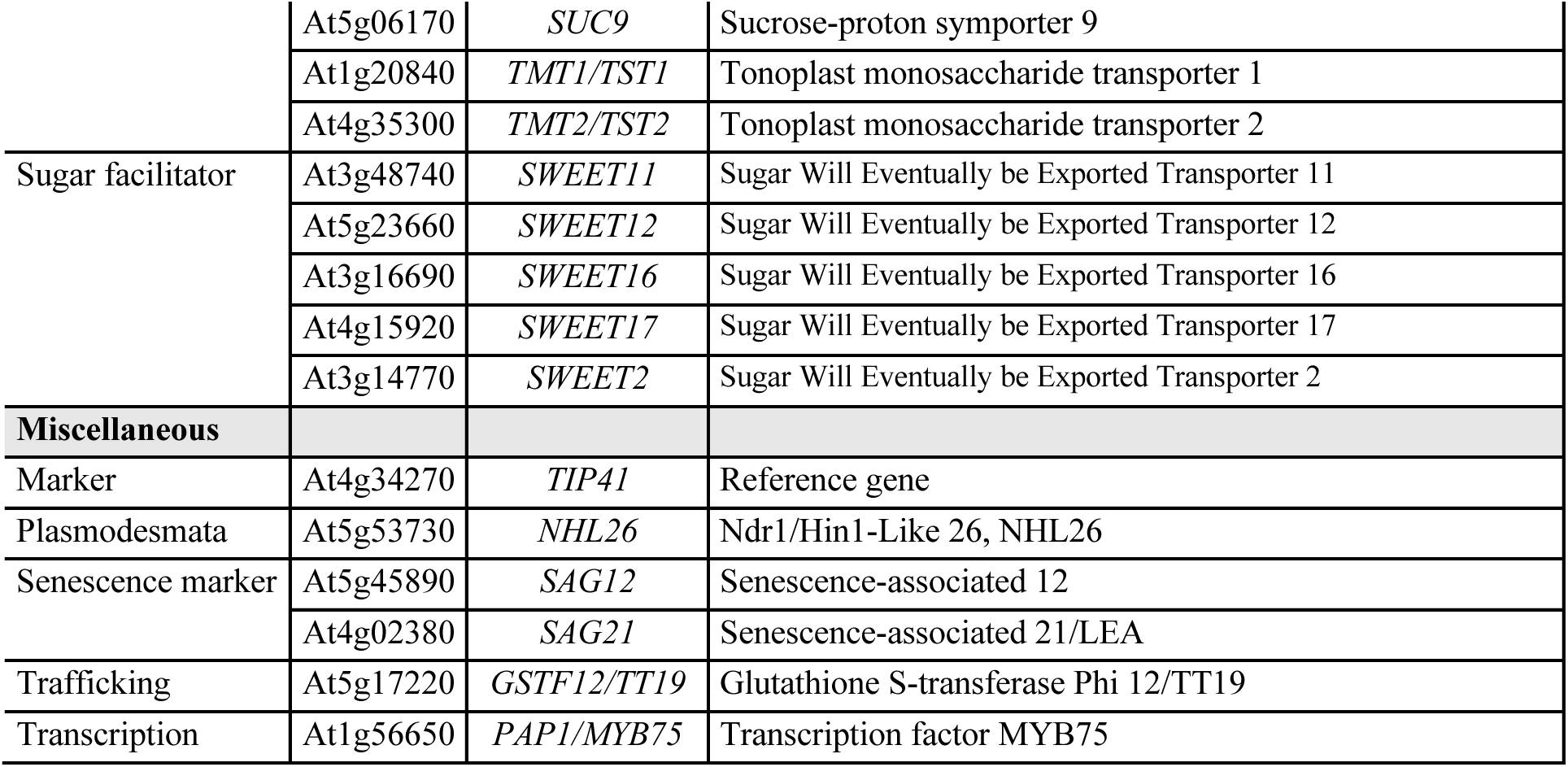
List of genes analyzed by RT-qPCR.

**Table S6.** Correlations (R_Pearson_) between gene expression and sugar accumulation in rosette leaves.

| Gene | Correlation | Pval | Gene | Correlation | Pval |
| --- | --- | --- | --- | --- | --- |
| *SUC1* | 0.7594 | <.0001 | *cwINV1* | 0.3872 | 0.0113 |
| *SWEET11* | 0.7550 | <.0001 | *cwINV6/FEH* | 0.6925 | <.0001 |
| *SWEET12* | 0.7795 | <.0001 | *FRK1* | 0.6635 | <.0001 |
| *G6PT2* | 0.8284 | <.0001 | *FRK2* | 0.4958 | 0.0008 |
| *APL3* | 0.7345 | <.0001 | *LHCB1* | -0.5347 | 0.0059 |

